# Increasing intraspecific plant chemical diversity at plot and plant level affects arthropod communities

**DOI:** 10.1101/2024.07.10.602874

**Authors:** Lina Ojeda-Prieto, Eliecer L. Moreno, Robin Heinen, Wolfgang W. Weisser

## Abstract

1. Plant chemistry mediates interactions between plants and their environment. While intraspecific chemodiversity at the plant level is well-studied, the effects of chemodiversity at the community level on arthropod interactions need more attention. We conducted a field experiment to test how intraspecific chemodiversity affects plant-arthropod interactions.

2. We manipulated plots of *Tanacetum vulgare* L., varying in chemotype richness and composition, and monitored four arthropod groups (herbivores, flower visitors, predators, and ants) over three seasons. We hypothesized that higher plot-level chemotype richness would enhance *occurrence* across all studied arthropod groups but have functional group-specific effects on *abundance*, resulting in reduced herbivore and ant abundance and increased flower visitor and predator abundance with increased chemotype richness.

3. Using mixed models, we found that increasing plot-level chemotype richness had a limited effect on most arthropod group occurrences but led to significant changes in abundance. Herbivore abundance decreased over time, and flower visitor abundance increased, while predatory arthropods and ants remained largely unaffected. Furthermore, we found that chemotype presence within a plot showed year-to-year variation in its effects, particularly on herbivores in which positive effects turned negative over time, and on flower visitors in which the presence of specific chemotypes positively affected their abundances. Predators and ants, on the other hand, showed weaker and more variable responses to specific chemotypes.

4. These results suggest that different arthropod groups may respond to plant chemicals through different ecological mechanisms. Our research underscores the role of plant chemical diversity in shaping insect communities and contributing to ecosystem dynamics.

## Resumen

1. La química de las plantas juega un papel clave en las interacciones con su entorno. Si bien la quimiodiversidad intraespecífica a nivel de planta ha sido ampliamente estudiada, sus efectos a niel de comunidad sobre las interacciones con artrópodos requieren más investigación. En este estudio, realizamos un experimento de campo para evaluar cómo la diversidad química dentro de una especie de planta afecta las interacciones con artrópodos.

2. Establecimos comunidades de *Tanacetum vulgare* L., en las que variamos la riqueza y composición de quimiotipos. Monitoreamos cuatro grupos de artrópodos-herbívoros, visitantes florales, depredadores y hormigas-durante tres temporadas. Hipotetizamos que una mayor riqueza de quimiotipos a nivel de parcela incrementaría la *ocurrencia* de todos los grupos de artrópodos, pero tendría efectos específicos sobre la *abundancia* de cada grupo funcional, reduciendo la abundancia de herbívoros y hormigas, y aumentando la de visitantes florales y depredadores.

3. Modelos mixtos revelaron que, aunque la riqueza de quimiotipos tuvo un efecto limitado sobre la ocurrencia de la mayoría de los grupos de artrópodos, tuvo efectos significativos en la abundancia. La abundancia de herbívoros disminuyó con el tiempo, mientras que la de los visitantes florales aumentó. En los artrópodos depredadores y hormigas no se encontraron efectos significativos. Adicionalmente, observamos que la presencia de ciertos quimiotipos en la comunidad tuvo efectos que variaron entre años: los herbívoros experimentaron un cambio de efectos positivos a negativos con el tiempo, mientras que los visitantes florales fueron consistentemente favorecidos por la presencia de ciertos quimiotipos. En contraste, los depredadores y hormigas mostraron respuestas más variables y menos pronunciadas a estos compuestos.

4. Nuestro estudio sugiere que distintos grupos de artrópodos responden de manera diferencia a las señales químicas de las plantas, probablemente debido a mecanismos ecológicos específicos para cada interacción. Estos resultados destacan el papel de la diversidad química de las plantas en la estructuración de las comunidades de artrópodos y su influencia en las dinámicas a nivel de ecosistema.

## Introduction

Biodiversity is recognized as a driver of ecosystem functioning and stability (Mora et al., 2014; Espinosa-García, 2022). Biodiversity studies have focused on species richness, examining how increasing plant community diversity affects ecosystem-level processes (Schulze & Mooney, 1992; Cardinale et al., 2006; Eisenhauer et al., 2011; Weisser et al., 2017). Plant communities with greater species richness are linked to a more diverse community of organisms (Ebeling et al., 2018), alterations in food web structure (Giling et al., 2019), and increased ecosystem process rates (Hertzog et al., 2017; Meyer et al., 2017). However, species-level diversity alone does not fully capture the complexity of ecological interactions. While species-level diversity has been well-studied, recent research explores biodiversity within species (Crutsinger et al., 2006; Koricheva & Hayes, 2018; Raffard et al., 2019; Noto & Hughes, 2020; Zerebecki & Hughes, 2023; Govaert et al., 2024; Konefal et al., 2024). Intraspecific diversity as a driver of key ecological processes and interactions within communities is increasingly recognized in biodiversity research (Kleine & Müller, 2011; Benedek et al., 2015; Bustos-Segura et al., 2017).

One component of within-species diversity is chemodiversity (Müller et al., 2020; Wetzel & Whitehead, 2020). Chemodiversity refers to the variety of chemical compounds produced by organisms (Whitehead et al., 2021; Walker et al., 2022). These compounds, including terpenoids, alkaloids, and phenolics, play diverse roles in plant defense (Hare, 2011; Wetzel et al., 2023), communication, and reproduction (Hartmann, 2007; Whitehead et al., 2022), potentially altering ecosystem dynamics like herbivory rates and pollinator preferences (Eisenring et al., 2021; Müller & Junker, 2022). Chemodiversity affects ecological interactions, impacting richness, diversity, and community structure (Dyer & Jeffrey, 2021; Fernandez-Conradi et al., 2022; Petrén et al., 2023). Studies have shown cascading chemodiversity effects within food webs (Kos et al., 2011; Bálint et al., 2016; Senft et al., 2019).

Despite growing evidence of the role of chemodiversity, particularly at the plant level, understanding how it operates at other ecological scales remain a key question. Intraspecific chemodiversity can be investigated at different scales, from individual plants to larger groups like plots or populations (Müller et al., 2020; Ojeda-Prieto et al., 2024). Research at the plant level has focused on chemodiversity effects on arthropod-plant interactions (Kos et al., 2011; Senft et al., 2017; Benedek et al., 2019). For instance, field observations show certain compounds influence herbivore preference and performance (Kleine & Müller, 2011; Clancy et al., 2018), as well as natural enemies’ behavior and reproductive success (Benedek et al., 2015; Zytynska et al., 2019). Manipulative studies reveal that plants with different chemical compositions differ in attractiveness to aphids (Neuhaus-Harr et al., 2024) and herbivore performance (Jakobs & Müller, 2018), reinforcing the role of chemodiversity in plant-arthropod interactions.

Larger-scale studies in chemodiversity are scarce and predominantly correlative. Studies have examined how local chemotype richness influences arthropod communities (Hauri et al., 2021; Espinosa-García, 2022), but many questions remain on the responses of different arthropod groups. Manipulative experiments, directly testing the consequences of a gradient of chemotype richness at the group level, show that increasing intraspecific chemodiversity of a plant community correlates with higher herbivore diversity, but lower plant damage (Bustos-Segura et al., 2017). By systematically manipulating **plot-level chemotype richness (CR)**, we aim to understand how chemodiversity at plant and plot level shapes arthropod abundance and occurrence. Due to their distinct ecological associations with plants, chemodiversity effects are expected to vary among arthropod groups. This complexity has not been fully addressed, highlighting the need for comprehensive experimental approaches on arthropod groups and their interactions with plants. Given the diversity of ecological roles of different arthropod groups, we expect chemodiversity effects to differ across herbivores, flower visitors, predators, and mutualists, as these interactions might be mediated by chemical cues and plant defenses (Petrén et al., 2023; Wetzel & Whitehead, 2020).

Different plant-associated arthropod groups may share different co-evolutionary histories with – and offer different selective pressures to - the plant that have likely shaped its chemical composition by selecting chemical profiles that serve optimal plant fitness (Dyer & Jeffrey, 2021; Schneider et al., 2021; Whitehead et al., 2021, 2022). Arthropod **abundance** is influenced by plant attractiveness and resource suitability, but it is also shaped by a complex interplay of factors, including predator-prey dynamics, trophic cascades, and habitat structure. However, conceptually, these relationships are best understood from a plant-centered perspective. Plant chemodiversity may be driven not just by herbivory pressure, but also by its effects on other interacting groups like pollinators and predators that suppress herbivores. For herbivores, plants may benefit from having a diverse bouquet of chemicals to deter herbivores and reduce their abundance and impact on the plant (Glassmire et al., 2019; Kariñho-Betancourt, 2020; Zu et al., 2020; Rabelo et al., 2023). For predatory arthropods, plants may benefit from their presence as predators may control herbivore populations (Barbosa et al., 2009; Turlings & Erb, 2018). Therefore, it is likely that more diverse blends of compounds may have evolved in part to attract predatory arthropods (De Boer et al., 2004, 2008; Clavijo McCormick et al., 2012). Similarly, many flower visitors play a crucial role in pollination (Jacobsen & Raguso, 2018; Dantas et al., 2020). Diverse chemical profiles may represent a reproductive advantage for the plants by attracting a wider range of pollinators. Plants may evolve specific compounds to attract more flower visitors, and more diverse profiles attract more visitors. For ants, the role of chemodiversity is likely to vary. Ants may have positive, negative, or no effects in different systems (Yamawo et al., 2021; Ramos et al., 2022). Given that many ants engage in mutualisms with herbivorous aphids, it may be beneficial for plants to evolve repellant compounds. On the other hand, if ants engage in protective mutualisms with plants, it may be beneficial for plants to evolve attractant compounds. Therefore, hypotheses regarding relationships between plant chemodiversity and ant abundance will likely depend on the role of the ants in the study system. Concerning the **occurrence** of arthropods, higher chemodiversity likely favors arthropod occurrence by increasing the likelihood that a specific compound can be used as a host recognition cue (Lampert et al., 2010; Dyer, 2018; Jacobsen & Raguso, 2018). As a chemical profile is, by definition, a result of cumulative selection processes that have taken place over evolutionary history, compounds may serve unique or dual purposes depending on the interactions. Understanding how plant chemodiversity impacts arthropod communities will provide insights into the ecological and evolutionary forces that shape biodiversity.

Despite extensive research on plant chemodiversity’s effects on arthropod communities, gaps remain. Studies often test only one arthropod group, focus on short-term effects, or compare high vs. low chemodiversity without examining a gradient. Our study addresses these gaps by manipulating the chemodiversity gradient and observing the responses of different arthropod groups over several seasons. We used *Tanacetum vulgare* L. (tansy), an herbaceous plant with high intraspecific variation in terpenoids, to investigate how variation in the number of tansy chemotypes in a plot affects associated arthropod communities. In a field experiment with 84 plots (six plants each), we manipulated **plot-level chemotype richness** (**-CR-**; i.e., 1, 2, 3, or 6 chemotypes present in a plot) and studied its effects on herbivores, flower visitors, predators, and ants (Fig. 1). We examined the effects of chemodiversity on arthropod occurrence and abundance at the plot level, recognizing that responses may differ across groups due to their distinct ecological roles and interactions with plants. Additionally, we investigated the effects of each of the six chemotypes on arthropod communities, anticipating differential responses based on compound attractiveness or repellence. We specifically tested the following hypotheses:

**Figure 1.**
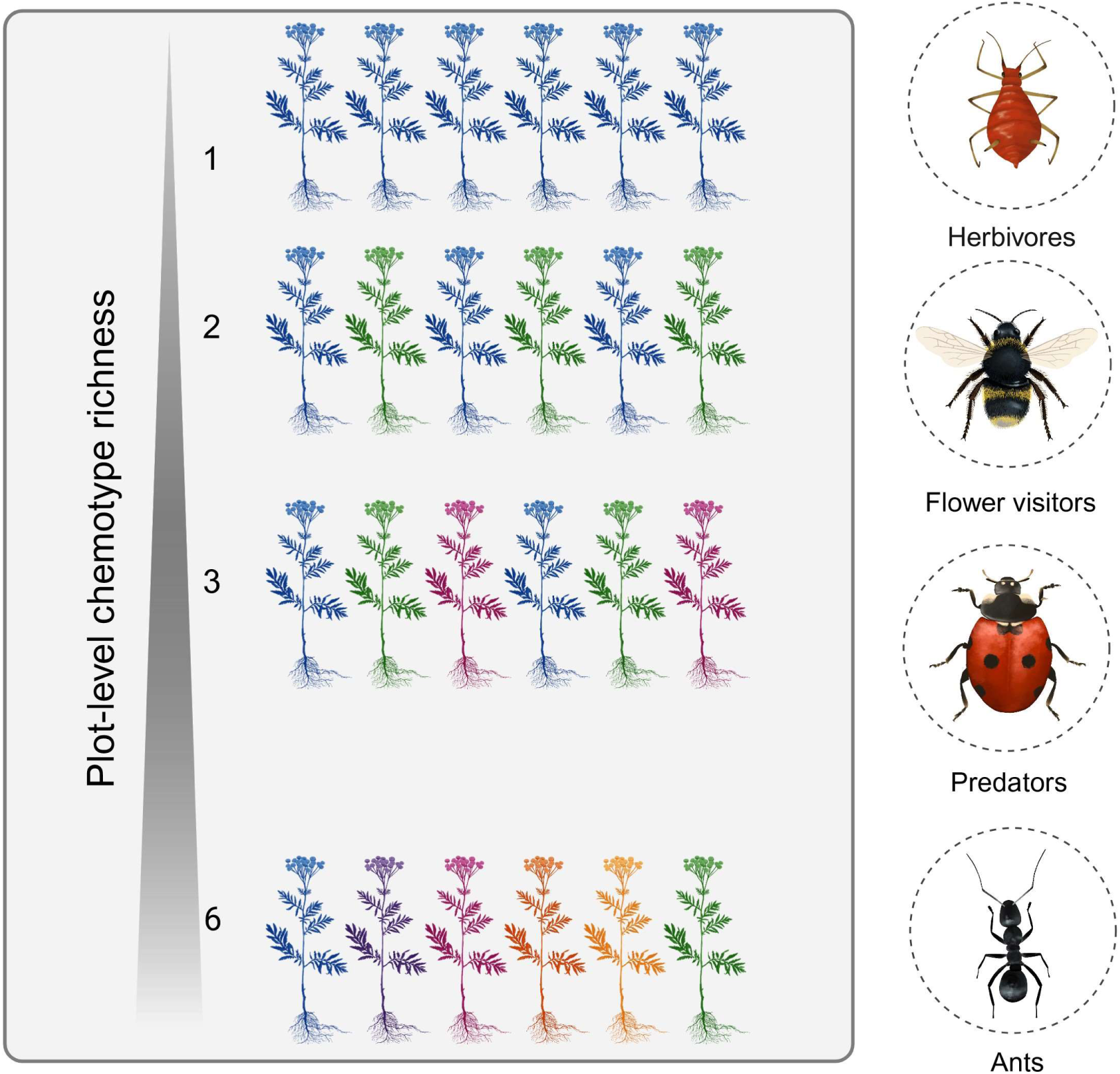
Influence of *Tanacetum vulgare* intraspecific chemodiversity on arthropod communities. The left panel illustrates the gradient of plot-level chemotype richness, with each color symbolizing a specific chemotype of *Tanacetum vulgare* in a six-plant community. From levels 1 to 6, the chemodiversity increases, indicating an increase in different chemotypes. The right panel presents the evaluated arthropod communities: herbivores, flower visitors, predators, and ants, which are expected to be influenced by intraspecific chemodiversity. Created with Biorender®. Agreement number: FE271K6HDD.

H1: Plots with greater chemotype richness will show increased arthropod occurrence across all trophic levels compared to plots with fewer chemotypes due to a wider range of available chemical cues that could lead to attraction of arthropod groups.

H2: Increased plot chemotype richness alters plant-arthropod associations, leading to differential effects on herbivores, flower visitors, predators, and mutualist abundances. We hypothesize that higher chemotype richness will reduce herbivore and ant abundances due to increased associational resistance, where chemically diverse plant communities reduce herbivore pressure. In contrast, flower visitor and predator abundance will increase with higher chemotype richness, as greater chemical complexity may increase resource attractiveness and prey availability.

H3: Individual plant chemotypes differentially influence plant-arthropod interactions based on their unique terpenoid profiles. We hypothesize that specific chemotypes will either attract or repel particular arthropod groups depending on their ecological roles. These different effects will shape the occurrence and abundance patterns of herbivores, predators, flower visitors, and ants.

## Materials and Methods

### Experimental design

Tansy chemotypes were characterized based on their leaf terpenoid compositions. We selected six chemotypes (’Athu-Bthu’, ‘Bthu-high’, ‘Bthu-low’, ‘Chrys-acet’, ‘Mixed-high’, and ‘Mixed-low‘) with three biological replicates each (Fig. S1-1). These chemotypes were propagated via stem cuttings and utilized in a field experiment at the Jena Experiment site in Germany in 2021. The experiment consisted of 84 1m^2^ plots, each containing six tansy plants (Table 1). The plots were designed to vary in chemotype richness levels ranging from 1 to 6 (Fig. S1-2), with 12, 30, 30, and 12 replicate plots, respectively, for each level. Each plot’s chemotype composition was pre-assigned, ensuring biological replication of each chemotype combination across the experiment. Plots were distributed equally among six randomized blocks, with specific numbers of plots allocated per chemotype richness level within each block (Ojeda-Prieto et al., 2024).

**Table 1.**
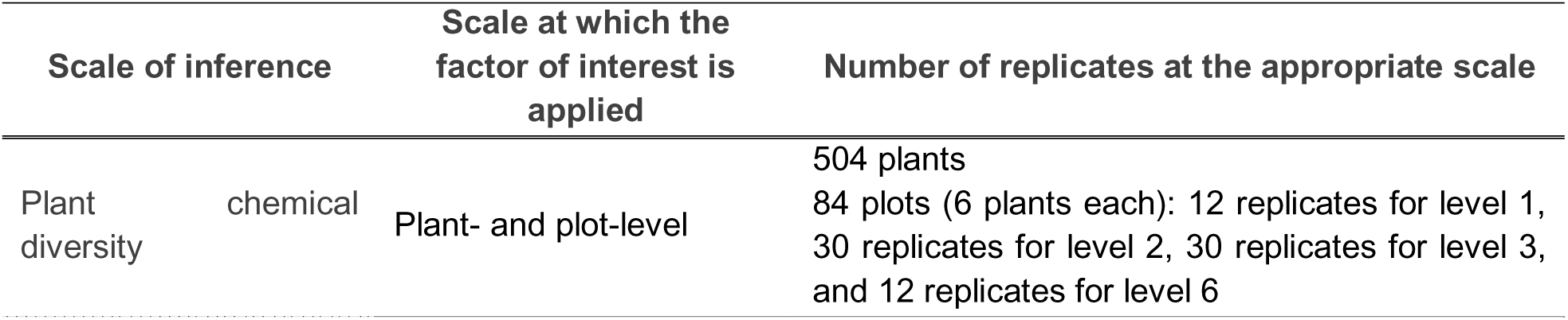
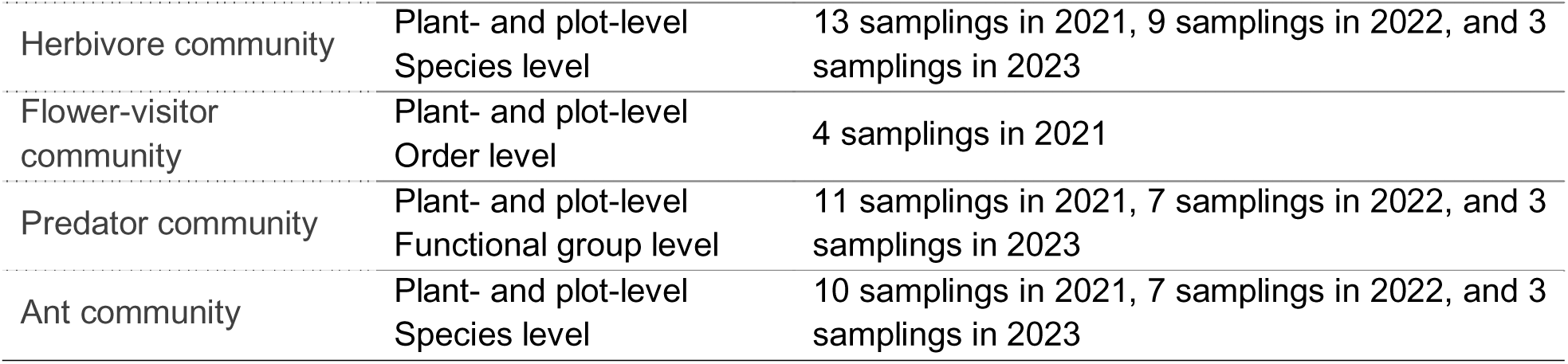
Information on the level of replication of the field experiment.

### Insect sampling

From 2021 to 2023, we assessed herbivores, flower visitors, predators, and ants (Table 1). Specific sampling dates are shown in Table S1-1. Each year, herbivore communities were monitored by counting individual aphids on the plants from their first appearance until nearly none were found. Although the focus was on aphids, other herbivores were scarce. Therefore, the generalization to herbivores here reflects predominantly aphids. Counts were conducted 13 times from June to August 2021, nine times from May to August 2022, and three times from June to August 2023. Each count, executed within a day between 7h and 19h, involved a meticulous scan of each plant, which started from the tip of each stem and proceeded to the end, examining both sides of each leaf. Identification of aphid species followed the key provided in Supplementary Information 1 (S1). When multiple aphid species were present on a plant, we recorded the total number of each. In cases of high aphid densities (>100), we estimated the total by counting aphids on 1 cm of stem or 1 cm^2^ of leaf three times in different stems and leaves for each plant, then multiplying the average by the estimated lengths of stems (in cm) or the estimated areas of leaves (in cm^2^) colonized by the aphids. We used plot-level occupancy and cumulative plot-level number of aphids to analyze occurrence and abundance.

Flower visitors were assessed by counting the flower heads of individual plants during the flower-visiting recording. Flower visitors were counted three times in July 2021 and once in August 2021. Surveys lasted three minutes per plot and were conducted in random block order from 9h to 15h under optimal weather conditions (no gusts of wind or precipitation). Flower visitors were identified at the order level. We used presence absence data to analyze the effects of CR and chemotype presence on occurrence. However, as CR and individual chemotypes influence reproductive traits in our system (Ojeda-Prieto et al., 2024), flower visitor abundance per flower head (visitation rate) assessed effects on abundance.

Predatory arthropods and ants on individual plants were recorded during herbivore counting for 2021 and 2022 and the day after in 2023. Predators were classified into taxa/feeding guilds (Araneae, Coleoptera, Dermaptera, Diptera, Hemiptera, and Hymenoptera), specifying aphid parasitoids and parasitized aphids (counted as mummies found). Ant presence on plants, but not abundance, was noted. Predator and ant occurrences and parasitized aphid abundance were used for analyses for the three seasons. In 2023, predator and ant abundances were also analyzed.

### Statistical analyses

Statistical analyses were conducted using R version 4.1.2 and RStudio 2023.09.1 (R Core Team, 2021). All statistics were performed at the plot level. The effects of chemotype presence and CR on the occurrence of herbivores, flower visitors, predators, and ants were assessed through binomial Generalized Linear Mixed-effect Models (GLMM) using the ‘*lme4*’ package (Bates et al., 2015). Linear Mixed-Effect Models (LMM) and Zero-Inflated Negative Binomial Models (ZINB), using the “*glmmTMB*” package (Brooks et al., 2017), modeled the effect on abundances. P-values were estimated by type II Wald-Chi-Squared tests using the Anova() function (‘*car*’ package; Fox & Weisberg, 2019). Data visualization used academic-licensed Biorender® and ‘*ggplot2’* (Wickham, 2009), *‘grid’* (R Core Team, 2021)*, ‘gridExtra’* (Auguie et al., 2017), *‘ggpubr’* (Kassambara, 2020), and *’DHARMa’* (Hartig, 2022) packages.

*Effects of chemotype richness and chemotype presence on arthropod taxa occurrence and abundance* Binomial GLMMs assessed the effect of CR and chemotype presence on occurrence. LMMs and ZINB tested the effects on abundance. Data were analyzed at the arthropod group and lower taxonomic levels (when indicated). Data were analyzed cumulatively for each year and for each independent sampling time. Detailed model conditions and transformations for each group are given in Supplementary Information 1 (S1).

## Results

During our sampling campaigns, we registered the following arthropod numbers: For 2021, 166.630 herbivores in total for 13 sampling dates (58.550 *Aphis fabae* (Scopoli, 1763), 22.691 *Brachycaudus cardui* (Linnaeus, 1758), 382 *Macrosiphoniella tanacetaria* (Kaltenbach, 1843), 62.055 *Metopeurum fuscoviride* (Stroyan, 1950), and 22.952 *Uroleucon tanaceti* (Linnaeus, 1758)), 974 flower visitors in four sampling dates (316 Coleoptera, 236 Diptera, 134 Hemiptera, 237 Hymenoptera, and 51 Lepidoptera), and 4.233 parasitized aphids in 10 sampling dates. For nine sampling dates in 2022, 11’171.066 herbivores (1.610 *A. fabae,* 3.321 *B. cardui*, 35.902 *Ma. tanacetaria*, 37.196 *Me. fuscoviride*, and >1.1 Mio *U. tanaceti*), and 341 parasitized aphids. For three sampling dates in 2023, 234.386 herbivores (249 *A. fabae,* 217 *B. cardui*, 0 *Ma. tanacetaria*, 9.535 *Me. fuscoviride*, and 224.385 *U. tanaceti*), 9.940 predators (1.573 Araneae, 4.770 Coleoptera, 508 Dermaptera, 1.032 Diptera, 531 Hemiptera, 5.309 Hymenoptera) and 3.783 ants (*Lasius niger* (Linnaeus, 1758)).

### Seasonal patterns in the herbivore community

Phloem feeders heavily dominated the free-living insect herbivore community in the tansy model system, whereas chewing herbivores were largely absent. We observed two generalist aphid species in our field study (*A. fabae* and *B. cardui*) and three tansy-specialist aphid species (*Ma. tanacetaria, Me. fuscoviride,* and *U. tanaceti)*. Our observations revealed the species-specific seasonal pattern of aphid populations. Populations grew exponentially during the warmer months of June and July and collapsed in August each year (Fig. S1-2). However, the timing and height of peak abundance varied among the species. The two generalist species (*A. fabae* and *B. cardui*) exhibited a similar pattern; they were the first to colonize the tansy plants and reached peak abundance in early July 2021 (Fig. S1-2a). However, they were much less abundant in 2022, with a peak occurring at the end of May. Conversely, the specialist species were less abundant in 2021, reaching their peak in mid-June for *Ma. tanacetaria* and in mid-July for *Me. fuscoviride* and *U. tanaceti.* In 2022, these species were more abundant, displacing the generalist species, and reached peak abundance in mid to late June (Fig. S1-2b). Generalist aphid species were even less common in the field in 2023, while *U. tanaceti* dominated the plots with increased abundance in July (Fig. S1-2c).

### Effects of chemotype richness and chemotype on herbivore occurrence and abundance

Our study first investigated the effect of CR on plot-level I) herbivore occurrence and II) herbivore abundance across samplings in each experimental year. All plots were occupied by at least one herbivore each year, but the herbivore species-specific occurrence in plots varied. For instance, *Me. fuscoviride* was present in all plots in 2021 (Table S1-2), for *U. tanaceti* this was the case in 2022 (Table S1-3) and 2023 (Table S1-4). The only significant effect of CR on aphid occurrence was observed for *Me. fuscoviride* in 2023, which decreased when CR increased (X^2^_1_ = 9.42, P = 0.002).

We then examined how specific tansy chemotypes influenced herbivore occurrence at the plot level using individual models for each chemotype (S1). Certain chemotypes were associated with higher or lower likelihoods of colonization, i.e., occurrence, and higher or lower herbivore abundance, depending on the year and herbivore species (Table S1-11). *A. fabae* occurrence in 2022 was higher when ‘Bthu-low’ (X^2^_1_ = 4.50, P = 0.034) or ‘Mixed-high’ was in the plot (X^2^_1_ = 6.68, P = 0.010), but lower when ‘Mixed-low’ was in the plot (X^2^_1_ = 4.89, P = 0.027). In 2023, it was lower when ‘Bthu-low’ (X^2^_1_ = 4.50, P = 0.034) or ‘Mixed-low’ (X^2^_1_ = 4.89, P = 0.027) was present and higher when ‘Mixed-high’ was present (X^2^_1_ = 6.68, P = 0.010). Certain chemotypes within the plot affected *B. cardui* occurrence in 2022; ‘Bthu-high’ increased *B. cardui* occurrence (X^2^_1_ = 4.16, P = 0.041), while ‘Bthu-low’ decreased it (X^2^_1_ = 6.42, P = 0.011). *Ma. tanacetaria* occurrence was not affected by any chemotype. *Me. fuscoviride* occurrence in 2023 decreased when ‘Athu-Bthu’ (X^2^_1_ = 5.23, P = 0.022) or ‘Mixed-high’ (X^2^_1_ = 9.42, P = 0.002) was present in the plot. *U. tanaceti* occurrence in 2021 decreased when ‘Mixed-high’ was present (X^2^_1_ = 7.47, P = 0.002).

Concerning abundance, there was no effect of CR on cumulative aphid abundance across all species (X^2^_1_ = 0.29, P = 0.588) nor within each of the five species in the first year (*A. fabae:* X^2^_1=_ 0.03, P = 0.870; *B. cardui:* X^2^_1_ = 1.51, P = 0.219; *Ma. tanacetaria:* X^2^_1=_ 1.13, P = 0.288; *Me. fuscoviride:* X^2^_1=_ 0.52, P = 0.471; *U. tanaceti:* X^2^_1=_ 0.38, P = 0.536; Fig. 3a, Table S1-2). However, increasing CR negatively affected herbivore abundance in the following years (2022: X^2^_1=_ 7.63, P = 0.006; 2023: X^2^_1_ = 4.39, P = 0.036; Fig. 2a, Tables S1-3, S1-4), mainly driven by the negative effect of CR on *U. tanaceti* populations in 2022 (X^2^_1_ = 5.93, p = 0.015) and on *U. tanaceti* (X^2^_1_ = 4.44, P = 0.035) and *B. cardui* abundances in 2023 (X^2^_1_ = 5.56, P = 0.018; Fig. 3a). Notably, *Ma. tanacetaria* aphids were not observed in the field in 2023 at all. Plot-level chemotypic diversity had little effect on whether a plot was occupied by aphids during the year, mainly because all plots were occupied. However, there were negative effects on the abundance of several herbivore species.

**Figure 2.**
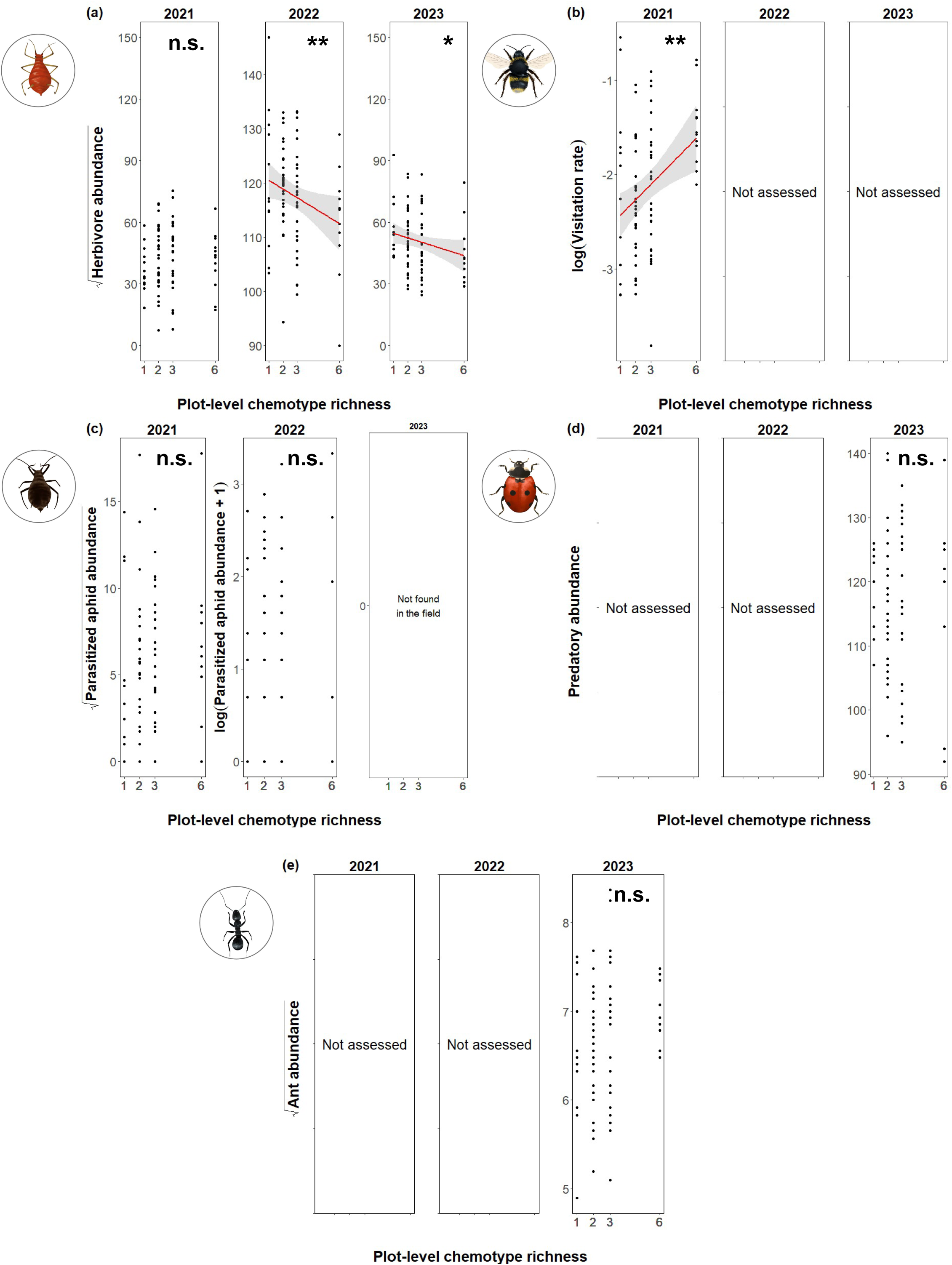
Effect of plot-level chemotype richness on the abundance of different arthropod communities over three years (2021–2023). **(a)** Effect on herbivore abundance, represented by five aphid species each year. **(b)** Effect on flower visitor abundance (i.e., visitation rate), calculated as the cumulative count of flower-visiting insects divided by the sum of flower heads counted during each assessment date. **(c)** Effect on parasitized aphid abundance (aphid mummies). **(d)** Effect on predator abundance. **(e)** Effect on ant abundance. Significance is indicated as follows: n.s. = not significant, * P < 0.05, ** P < 0.01, and *** P < 0.001. Degrees of freedom, Wald’s Chi-square statistics, and p-values are reported in Tables S1 – S8.

**Figure 3.**
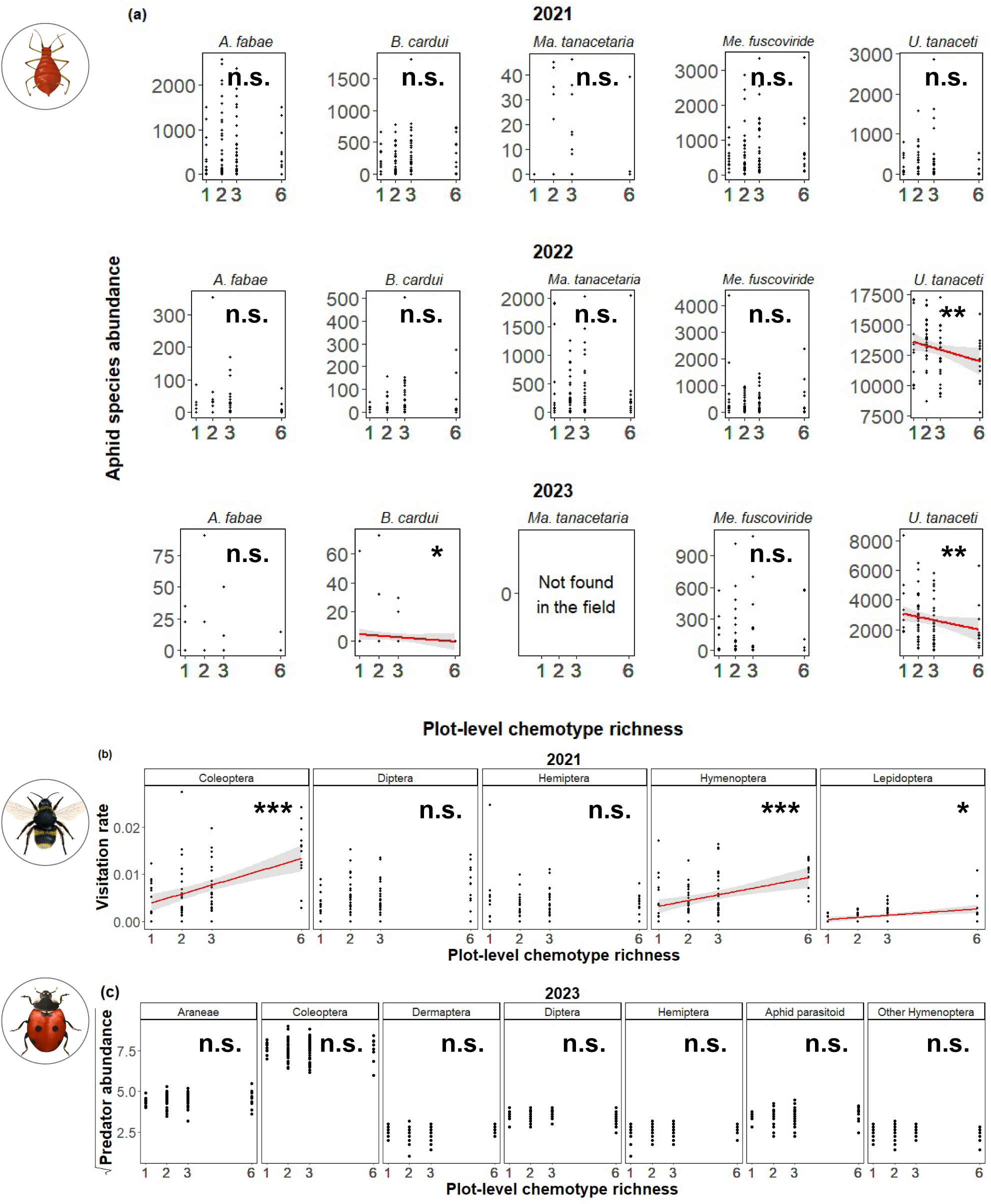
Effect of plot-level chemotype richness on specific group abundances within each guild of arthropod communities over three years (2021–2023). **(a)** Effect on herbivore abundance of *Aphis fabae*, *Brachycaudus cardui*, *Macrosiphoniella tanacetaria*, *Metopeurum fuscoviride*, and *Uroleucon tanaceti* each year. **(b)** Effect on flower visitor abundance of Coleoptera, Diptera, Hemiptera, Hymenoptera, and Lepidoptera. **(c)** Effect on predator abundance of Araneae, Coleoptera, Dermaptera, Diptera, Hemiptera, Aphid parasitoids, and Other Hymenoptera for 2023. Significance is indicated as follows: n.s. = not significant, * P < 0.05, ** P < 0.01, and *** P < 0.001. Degrees of freedom, Wald’s Chi-square statistics, and p-values are reported in Tables S1 – S7.

If there was an effect of a specific chemotype present in a plot on herbivore abundance in the year 2021, it was always positive. However, all significant effects of particular chemotypes observed in 2022 or 2023 were negative (Table S1-11). Herbivore abundance was lower in 2022 in either ‘Chrys-acet’ or ‘Mixed-high’ presence and lower in 2023 when ‘Athu-Bthu’ or ‘Mixed-low’ was in the plot (Tables S1-3 and S1-4). When investigating species-specific effects, we found that *A. fabae* abundance in 2021 increased when ‘Mixed-high’ was present (X^2^_1_ = 6.77, P = 0.009), *B. cardui* when either ‘Bthu-high’ (X^2^_1_ = 3.97, P = 0.046) or ‘Chrys-acet’ were present (X^2^_1_ = 6.16, P = 0.013), and *U. tanaceti* when ‘Bthu-high’ was in the plot (X^2^_1_ = 5.98, P = 0.014). Lower *A. fabae* abundance was observed in 2022 when ‘Bthu-low’ was in the plot (X^2^_1_ = 3.91, P = 0.048) and in 2023 when ‘Athu-Bthu’ (X^2^_1_ = 7.39, P = 0.006) or ‘Chrys-acet’ (X^2^_1_ = 19.14, P < 0.001) were part of the community. *Me. fuscoviride* abundance in 2022 was negatively associated with ‘Bthu-low’ (X^2^_1_ = 5.85, P = 0.016), and *U. tanaceti* abundance in 2022 was negatively associated with either ‘Chrys-acet’ (X^2^_1_ = 12.77, P < 0.001) or ‘Mixed-high’ (X^2^_1_ = 5.58, P = 0.018). In 2023, *B. cardui* was lower when ‘Athu-Bthu’, ‘Bthu-high’, or ‘Mixed-high’ was present (Athu-Bthu: X^2^_1_ = 4.42, P = 0.035; Bthu-high: X^2^_1_ = 40.71, P < 0.001; Mixed-high: X^2^_1_ = 79.26, P < 0.001), and *U. tanaceti* abundance was negatively associated to the presence of ‘Athu-Bthu’ (X^2^_1_ = 5.08, P = 0.024) or ‘Mixed-low’ (X^2^_1_ = 3.92, P = 0.048). The effects of specific chemotypes on herbivore abundance varied across different years, with positive impacts observed in 2021 and predominantly negative impacts in 2022 and 2023.

### Effects of chemotype richness and chemotype identity on flower visitors

We first analyzed the effect of CR on flower visitor occurrence. Higher CR increased the occurrence of Lepidoptera, i.e., adult butterflies (X^2^_1_ = 3.87, P = 0.049), but not other taxa. Hymenoptera and Lepidoptera flower visitor occurrences increased when ‘Bthu-low’ and ‘Athu-Bthu’ were in the plot, respectively (Tables S1-5, S1-11).

We evaluated the effect of CR on the visitation rate of flower visitors. We found a positive effect on all orders (X^2^_1_ = 10.71, P = 0.001, Fig. 2b), and specifically for Coleoptera (X^2^_1_ = 16.78, P < 0.001), Hymenoptera (X^2^_1_ = 11.56, P < 0.001), and Lepidoptera (X^2^_1_ = 8.80, P = 0.003, Fig. 3b). Concerning the presence of particular chemotypes within plots, ‘Chrys-acet’ (X^2^_1_ = 5.98, P = 0.014) and ‘Mixed-high’ (X^2^_1_ = 6.95, P = 0.008) positively affected the visitation rate across all the orders. While Diptera and Hemiptera abundances were not influenced by the presence/absence of any chemotypes, Coleoptera was influenced by the presence of ‘Bthu-low’ and ‘Chrys-acet’ (Table S1-5); Hymenoptera by ‘Bthu-low’ (X^2^_1_ = 7.58, P = 0.006), and Lepidoptera by ‘Athu-Bthu’ (X^2^_1_ = 7.53, P = 0.006, Table S1-12).

Analysis of the effect on the total number of individuals (not the visitation rate) is in Table S1-6. We found positive effects on total number of individuals of all orders (X^2^_1_ = 10.71, P = 0.001), Coleoptera (X^2^_1_ = 16.78, P < 0.001), Hymenoptera (X^2^_1_ = 11.56, P < 0.001), and Lepidoptera (X^2^_1_ = 8.80, P = 0.003), as found by the analysis of CR on visitation rate. However, when using the total number as the response variable, particular chemotypes had different effects on different groups. The abundance of all orders was affected by the presence of any of the chemotypes except for ‘Chrys-acet’, ‘Athu-Bthu’ affected the number of flower visitors belonging to Coleoptera and Lepidoptera, ‘Bthu-low’ to Coleoptera and Hemiptera, and ‘Bthu-high’ and ‘Mixed-low’ to Coleoptera (Table S1-6).

### Effects of chemotype richness and chemotype on predators and ants

Predatory arthropods occurred in all plots in all years (Tables S1-7 to S1-9), and ants (*L. niger*) occurred in all plots in 2023 (Table S1-10). The only effect of CR on predator taxa/feeding guild occurrence was observed in predatory Hemiptera in 2022 (X^2^_1_ = 6.54, P = 0.010), which increased with CR. For chemotype identity effects, the occurrence of parasitized aphids in 2021 decreased when ‘Mixed-low’ was in the plot and in 2022 when ‘Chrys-acet’ was present (2021: X^2^_1_ = 4.71, P = 0.030; 2022: X^2^_1_ = 5.55, P = 0.018, Tables S1-7, S1-8).

Increasing CR did not increase predator abundance (overall-Fig. 2c-nor by taxa/feeding guild-Fig. 3c-). However, there was a positive effect of ‘Chrys-acet’ presence on the number of parasitized aphids in 2021 (X^2^_1_ = 6.30, P = 0.012, Table S1-7) and ant abundance in 2023 (X^2^_1_ = 6.58, P = 0.010, Table S1-8), but a negative effect on total predator abundance in 2023 (X^2^_1_ = 4.16, P = 0.041) and on abundance of predatory beetles in 2023 (Coleoptera: X^2^_1_ = 3.85, P = 0.050, Table S1-9). A positive effect on Dermaptera abundance in 2023 was found when ‘Athu-Bthu’ was present (X^2^_1_ = 8.51, P = 0.004).

### Temporal effects

Analyses at independent sampling times, available in Supplementary Information 1 (S1), revealed that while relationships between chemotype richness and the occurrence of various ecological groups were generally weak, the temporal patterns in abundance were clear. Effect on abundance showed a shift in slope from positive to negative for herbivores over time (particularly in specialist aphids); while it remained mostly positive relationships for flower visitors, ants, and predators (Supplementary Information 2).

## Discussion

Intraspecific chemodiversity is a key driver of interaction variation within plant species. Most of our understanding comes from observational and correlative studies, with controlled studies in natural or semi-natural settings limited due to the challenges of manipulating plant chemistry. In this study, we manipulated plant chemical diversity at the plot level by selecting plant chemotypes with varying chemical blends and manipulating the number of chemotypes within plots (i.e., **plot-level chemotype richness-CR-**). Using tansy (*Tanacetum vulgare* L.), we explored the effect of chemodiversity on natural arthropod communities. Our study provides evidence that intraspecific chemodiversity at the plot level influences arthropod abundance and occurrence. We showed that CR had negative effects on herbivores but positive effects on flower visitors, with minimal effects on predators or ants. This study also shows the strong effects of chemotype identity, i.e., the composition of the plots, on plant-arthropod interactions at the plot level, an effect also often found in biodiversity experiments (e.g., Scherber et al., 2010). Our study thus highlights the role of individual chemotypes in shaping interactions, contributing to our understanding of the ecological significance of chemical diversity in plants.

### Effects of chemotype richness on the occurrence of arthropod groups

Contrary to our expectations, CR had weak overall effects on the occurrence of most arthropod groups **(H1)**. While some significant relationships between CR and specific taxa occurrence were observed, they were often not strong. One reason may be that many arthropod taxa either occurred in all plots or were absent during certain samplings, limiting the variation needed to detect strong patterns. Another reason is that, for instance, specialized herbivore populations build up over the season before they decline, and in years when aphids are abundant, total population sizes are very high, and almost all plants are infested with each species at one point in time. We observed significant positive relationships for polyphagous aphid species such as *A. fabae* and *B. cardui* in the individual samplings during the first sampling year. These species only feed on tansy for several weeks, before they are replaced by the more abundant monophagous species. Significant negative relationships were observed for monophagous species, particularly in *Me. fuscoviride* in 2023, suggesting that polyphagous and monophagous species may respond differently to chemical cues in their host-seeking behavior (Gripenberg et al., 2010; Benedek et al., 2015; Ziaja & Müller, 2023). It also emphasizes the strong shift from polyphagous to monophagous herbivores in the system over the season and over the years. It could be that during the establishment phase, polyphagous herbivores were more abundant in the surrounding grassland and that monophagous species required time to colonize from the nearest tansy hosts. Specialized tansy aphids overwinter on or next to the host plant (Liu et al., 2017; Loxdale, Massonnet, et al., 2011; Loxdale, Schöfl, et al., 2011); hence, once the field plots are colonized, local overwintering and early colonization in the season by the specialized aphids are possible.

For flower-visiting taxa, however, we observed consistent positive relationships between CR and occurrence, indicating the role of specialized chemistry in these mutualists (Silva et al., 2018; Farré-Armengol et al., 2020; Sasidharan et al., 2023). These findings suggest that plant chemistry differentially affects arthropod groups based on their interaction with the plant. The lack of strong effects on predator and ant occurrences suggests that plant chemistry significantly may be more important for directly interacting organisms (e.g., herbivores and flower visitors) than for organisms that interact indirectly with the plant, such as predators and ants (Ali et al., 2023).

### Effects of chemotype richness on the abundance of arthropod taxa

We found strong support for CR affecting the abundance of herbivores and flower-visiting arthropods but not predatory arthropods and ants, partially in line with **(H2)**. Higher CR plots supported fewer herbivores in 2022 and 2023, although this relationship varied between aphid species. In general, the effects of CR on herbivore abundance were driven by its effects on the dominant monophagous aphid species, *U. tanaceti*, for both years and *B. cardui* for 2023. An interesting observation is that the slopes between CR and herbivore abundance in the individual sampling points, although not always significant, weres consistently positive in 2021 but turned increasingly negative over the season and across subsequent years. This shift towards monophagous aphids over time may emphasize the negative effect of CR on herbivore abundance. Since *U. tanaceti* had not yet successfully colonized the experiment in the first year, and polyphagous species were more common, this may explain the lack of CR effects in the first year. Monophagous herbivores may be more strongly affected by chemical defenses in high CR (Ziaja & Müller, 2023) in concordance with the neighborhood-mediated associational resistance hypothesis (Randlkofer et al., 2010).

Higher CR plots had more flower visitors, with significant positive effects observed for Coleoptera, Hymenoptera, and Lepidoptera but not for Diptera and Hemiptera, suggesting differential effects of CR on flower visitors (Sasidharan et al., 2023). Chemotypes also exhibit variations in their phenology, influenced by the chemical diversity of their growth environment (Ojeda-Prieto et al., 2024). These variations can extend the flowering period within plots containing more chemotypes, contributing to higher occurrence and abundance of flower visitors.

Our study revealed that plot-level chemotype richness did not affect predator or ant abundance. The reasons underlying these findings remain unclear, but additional factors, such as prey availability, which could be influenced by herbivore-host chemical composition (Benedek et al., 2019), or alternative foraging strategies (Wäschke et al., 2013), may contribute more strongly to these relationships than chemodiversity. Further research is warranted to elucidate the interplay between these factors and their collective impact on predator and ant abundance in response to chemodiversity.

### Effects of chemotype presence on arthropod communities

We found chemotype-specific impacts on plot-level occurrence and abundance across arthropod taxa, in line with our hypothesis **(H3)**. However, these patterns were rather variable across years. Herbivore occurrence data showed few chemotype-specific effects in 2021 – the first season after establishment. In subsequent years, both positive and negative effects of different chemotypes on herbivore species occurrence. This suggests that most aphids seem to respond to plant-chemical cues as expected (Mehrparvar et al., 2014; Clancy et al., 2018). Herbivore abundance data show a clear pattern, where in 2021, chemotype effects were typically positive, which may suggest that colonizers were using specific terpenoid cues to locate the plants initially. Interestingly, in later years, chemotype-specific effects, when significant, were consistently negative, suggesting that chemotypes may form a mosaic that partly regulates aphid densities, aligning with the idea that specialized chemistry evolved for defense (Fox, 1981; Wolf et al., 2011). This is further reinforced by the overall neutral-to-negative effect of chemotypes on herbivore abundance.

For flower-visiting arthropods, chemotype-specific patterns were most common in abundance data. Most flower visitors are highly mobile and do not linger in the same plot for long (Burdon et al., 2020; Knauer et al., 2021), implying that specific chemotypes attract higher number of individuals. Significant chemotypes positively affected flower visitors, with low-concentration B-thujone chemotypes having the most robust and consistent effects across taxa, partially aligning with previous results (Sasidharan et al., 2023). Mixed chemotypes showed few effects, while high-concentration B-thujone chemotype had no effect on flower visitors. Though B-thujone can signal nectar or pollen rewards (Metcalf & Kogan, 1987), its effects depend on the overall blend of volatile compounds and the preferences of local flower-visiting species. We found stronger effects of chemotype identity on the total number of individuals than on the visitation rate (Tables S1-5 and S1-6). As CR and individual chemotypes have been shown to influence reproductive traits in previous studies (Ojeda-Prieto et al., 2024), it is perhaps not surprising that effects of CR on total number of flower visitors were more pronounced than on visitation rates that were corrected for the number of reproductive stalks. Nevertheless, these results suggest that the direct CR effect is also robust and that the effect of individual chemotypes is also found when taking these reproductive traits into account.

Few, and only weak chemotype-specific effects were found for predators and ants, suggesting that these groups may be influenced by different mechanisms, including modified behaviors, such as predation activity or aphid-tending in ants (Senft et al., 2017; Mehrparvar et al., 2018; Hauri et al., 2022).

### Future perspectives for chemodiversity research

Exploring several key areas of chemodiversity will deepen our understanding. Scaling up experiments to entire landscapes will elucidate how chemodiversity influences ecosystems and biodiversity across diverse habitats. Assessing its effect on plant-insect interactions should include a focus on ecosystem functions like predation rates and herbivory impacts. Studying how chemodiversity influences herbivore behavior, particularly the responses of both generalist and specialist species, will offer insights with potential applications in pest management. Further exploring the interplay between plant chemistry and arthropod communities will advance our understanding of ecosystem functioning and guide future conservation and management practices.

### Conclusion

Previous studies have focused on intraspecific variability in ecosystem functioning. Our research demonstrates that plot-level chemotypic diversity impacts arthropod colonization patterns. Our results emphasize the role of intraspecific chemodiversity in shaping arthropod communities on plants. Specifically, we show that plots with higher intraspecific chemodiversity accumulate fewer herbivores and attract more flower visitors. The presence of specific chemotypes within a plot exerted highly variable impacts on abundance but also on occurrence patterns of various arthropod groups, suggesting that specific chemotypes, and potentially their specific chemical components, may drive the presence or absence of particular arthropod taxa.

## Notes

### Competing Interest Statement

The authors have declared no competing interest.

### Summary of Updates

Authors may wish to update the online version. No significant differences are found between this version and previous versions of the manuscript.

