## Supplementary material for "Increasing intraspecific plant chemical diversity at plot and plant level affects arthropod communities": S1

***New Phytologist* Supporting Information 1 (S1)**

**Article title:** Increasing intraspecific plant chemical diversity at plot and plant level affects herbivorous, predatory, and pollinating arthropod communities

**Authors:** Lina Ojeda-Prieto, Eliecer L. Moreno, Robin Heinen, Wolfgang W. Weisser

**Article acceptance date:**

The following Supporting Information is available for this article:

### 1. Supplementary methods

#### *1.1 Field design*

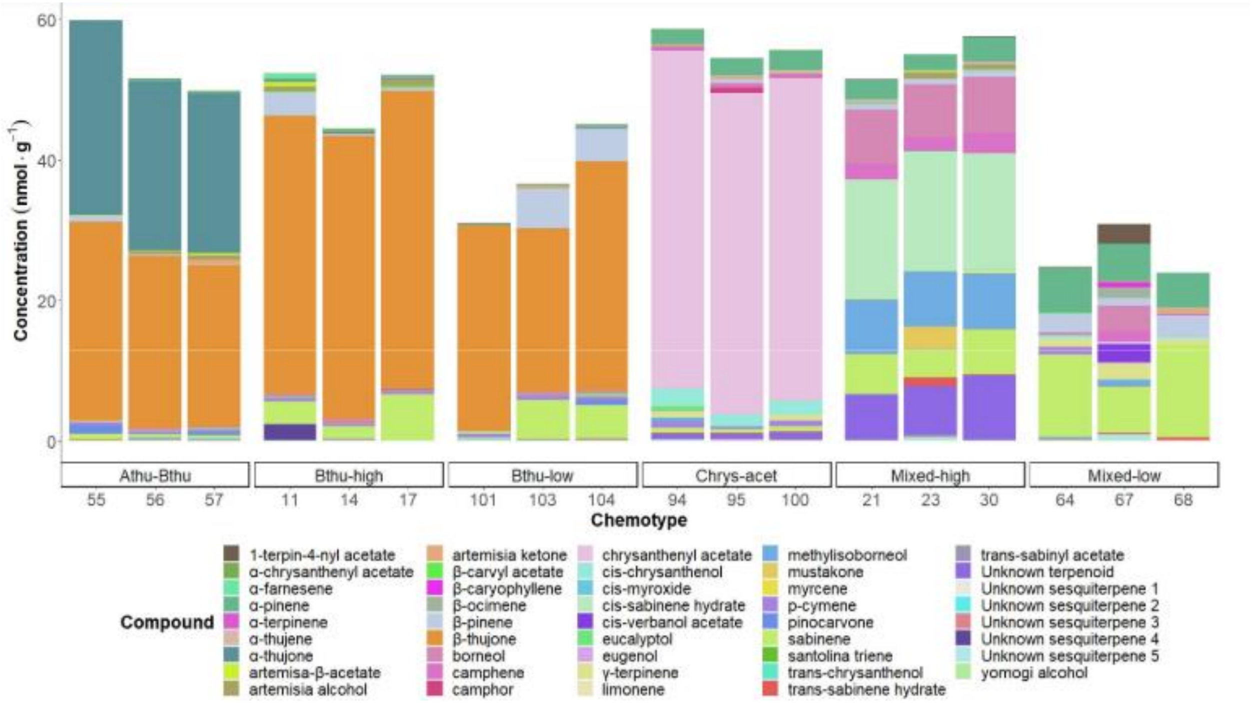

**Figure S1-1.** Stacked bar charts, adapted from Neuhaus-Harr et al., 2024 and reproduced from Ojeda-Prieto et al., 2024, display the approximate concentrations of terpenoid compounds (nmol g-1) extracted from leaf samples of 18 selected *Tanacetum vulgare* plants. These plants are organized by chemotype.

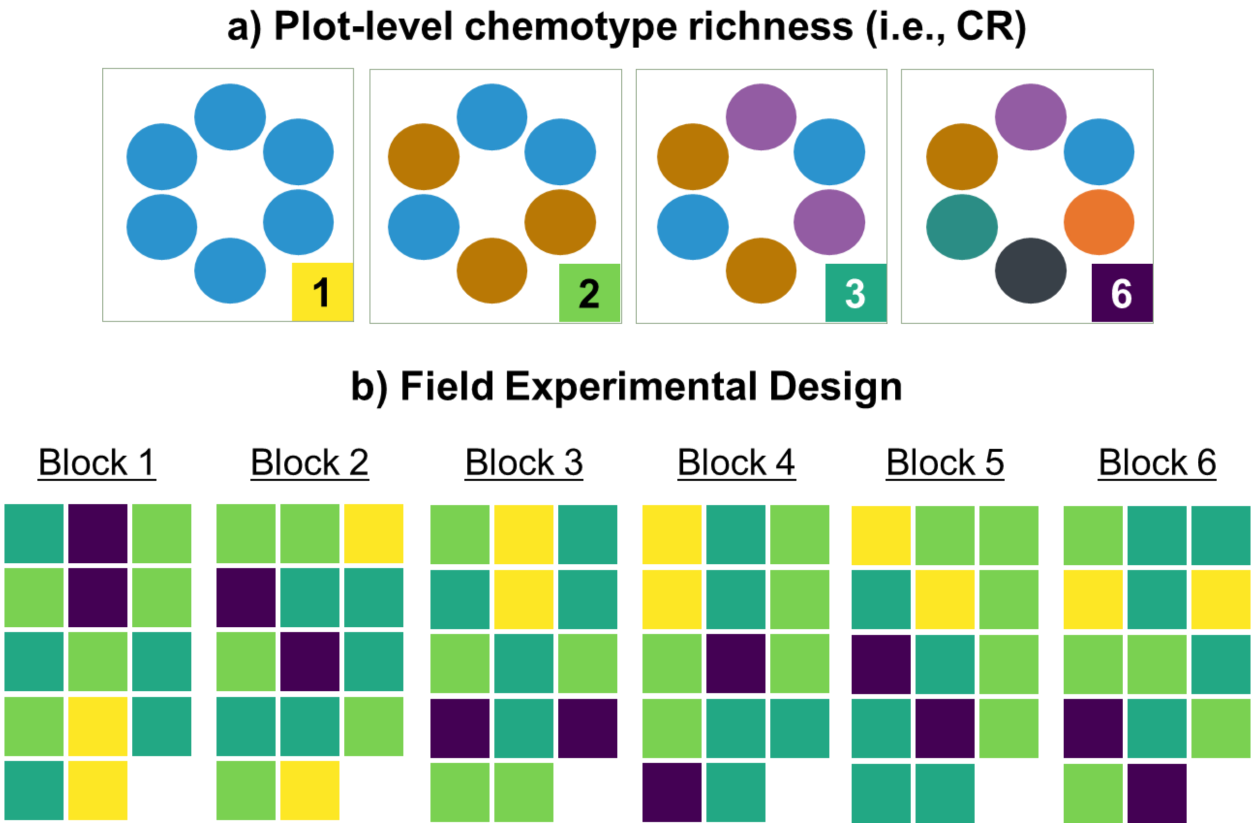

**Figure S1-2. (a)** Plot-level chemotype richness, which refers to the number of different chemotypes in a single plot, ranged from 1 to 6. **(b)** Block design. Section (a) displays the plot-level chemotype richness color scheme: yellow for 1, light green for 2, dark green for 3, and purple for 6. There were 84 plots in total, distributed equally across six randomized blocks. Each block contained 14 plots: two plots with chemotype richness level 1, five plots with level 2, five plots with level 3, and two plots with level 6.

### *1.2 Insect sampling*

*1.2.1 Sampling dates*

**Table S1-1.** Sampling dates in 2021, 2022, and 2023, specifying calendar week, specific date, and sampled arthropod group (herbivore, flower visitors, predators, and ants). Total counts across the three years are 25 for herbivores, 4 for flower visitors, 21 for predators, and 20 for ants.

| Year | Calendar week | Date | Sampled arthropod group |
|---|---|---|---|
| 2021 | 24 | Jun 07 | Herbivores |
| | 24 | Jun 11 | Herbivores |
| | 25 | Jun 14 | Herbivores, Predators, Ants |
| | 25 | Jun 18 | Herbivores, Predators, Ants |
| | 26 | Jun 21 | Herbivores, Predators, Ants |
| | 26 | Jun 25 | Herbivores, Predators, Ants |
| | 27 | Jun 28 | Herbivores, Predators, Ants |
| | 27 | Jul 02 | Herbivores, Predators, Ants |
| | 27 | Jul 03 | Flower visitors |
| | 28 | Jul 08 | Herbivores, Predators, Ants |
| | 28 | Jul 09 | Flower visitors |
| | 29 | Jul 15 | Herbivores, Predators, Ants |
| | 29 | Jul 16 | Flower visitors |
| | 30 | Jul 22 | Herbivores, Predators, Ants |
| | 31 | Jul 29 | Herbivores, Predators, Ants |
| | 32 | Aug 04 | Flower visitors |
| | 33 | Aug 10 | Herbivores, Predators |
| 2022 | 20 | May 14 | Herbivores, Predators, Ants |
| | 22 | May 25 | Herbivores, Predators, Ants |
| | 23 | May 31 | Herbivores, Predators, Ants |
| | 24 | Jun 07 | Herbivores, Predators, Ants |
| | 25 | Jun 15 | Herbivores, Predators, Ants |
| | 26 | Jun 21 | Herbivores, Predators, Ants |
| | 27 | Jun 27 | Herbivores, Predators, Ants |
| | 29 | Jul 13 | Herbivores |
| | 32 | Aug 03 | Herbivores |
| 2023 | 26 | Jun 20 | Herbivores |
| | 26 | Jun 21 | Predators, Ants |
| | 31 | Jul 25 | Herbivores |
| | 31 | Jul 26 | Predators, Ants |
| | 36 | Aug 29 | Herbivores |
| | 36 | Aug 30 | Predators, Ants |

*1.2.2 Aphid species identification key*

The following aphid species identification key is an English version based on Thieme & Müller (2011). All aphid species listed here are associated with *Tanacetum vulgare* L. plants. The identification key was complemented with figures, a summary of the main characteristics of each genus, and a glossary of terms.

**Common species on tansy:** *Aphis fabae, A. vandergooti, Brachycaudus cardui, B. helichrysi, Coloradoa tanacetina,* ***Uroleucon tanaceti****,* ***Metopeurum fuscoviride, Macrosiphionella millefolii, M. tanacetaria****, Uroleucon tanaceti.*

**1.** Cauda not longer than wide, semicircular, triangular or pentagonal......... ***Brachycaudus*** **(2)**

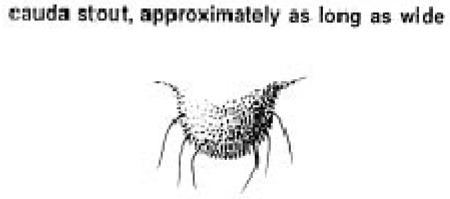

(Miller & Stoetzel, 1997)

**1*.** Cauda longer than broad, tongue- or finger-shaped or acutely triangular………. ***Aphidinae*, *Macrosiphoninae*** **(3)**

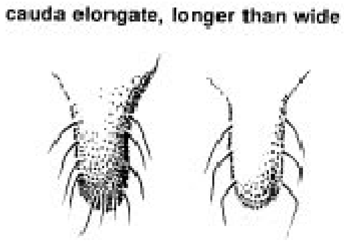

(Miller & Stoetzel, 1997)

**2.** Cornicles are shorter than the distance between the bases of the antennae (the terminal process of the last antennal segment is three or more times the basal part). Tip of proboscis exceeding the coxa of the third pair. Immatures are light green, sometimes reddish or orange. Body broadly rounded. 1.6-2.5 mm. Dorsal abdomen variably pigmented (dark sclerotic shield…………………………………………………….……… ***Brachycaudus cardui*** **(L.)**

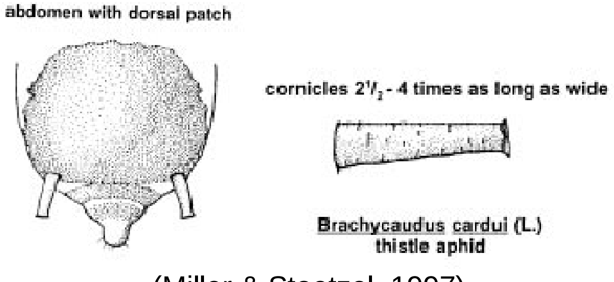

(Miller & Stoetzel, 1997)

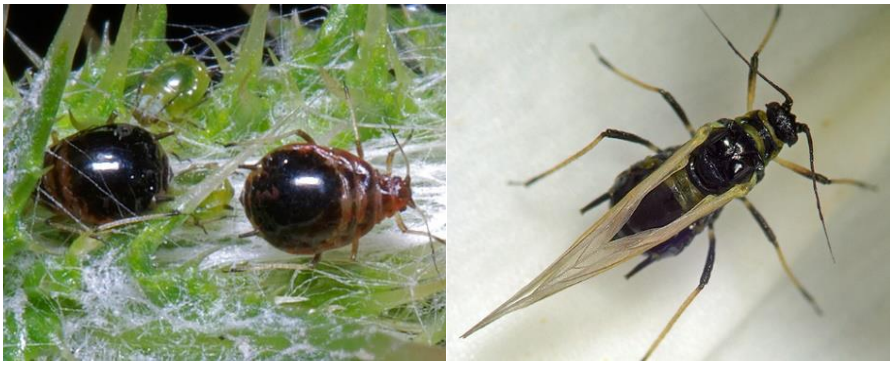

(InfluentialPoints, 2021)

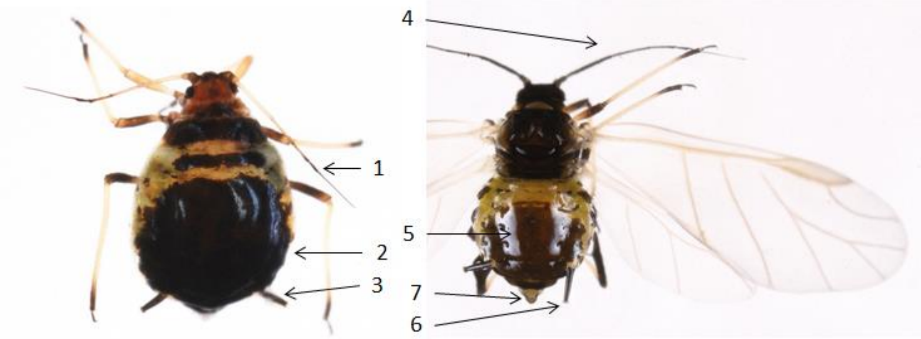

(Hullé et al., 2020)

**2*. Cornicles** are as long as or slightly longer than the distance between the bases of the antennae. Cornicles of unwinged colorless or only weakly tanned. The upper side of unwinged aphids (except fundatrix) is unpigmented. Body coloration is generally light green, sometimes yellowish, pink to white (without dorsal patch). Winged: Yellowish green with a large brown dorsal patch. Antennae are shorter than the body and have dusky tips. Siphunculi are pale, tapered, and short (0.8-2.0 times the length of the cauda) 1.2-2.2 mm … ***Brachycaudus helichrysi* (Kaltenbach)**

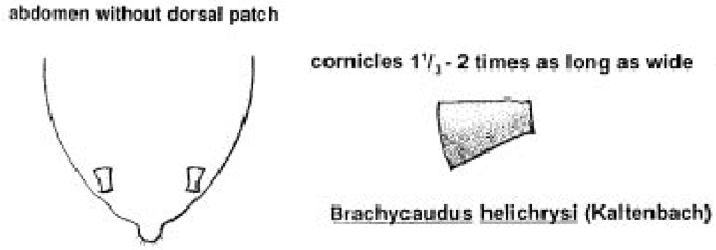

(Miller & Stoetzel, 1997)

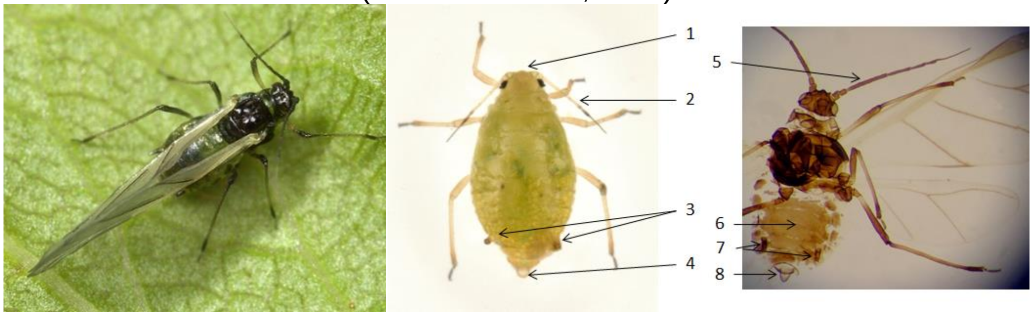

(Hullé et al., 2020; InfluentialPoints, 2021)

*Brachycaudus*

- Small to medium-sized oval aphids.
- Low lateral prominences.
- Antennae are shorter than the body.
- Siphunculi short to moderate in length.
- Cauda is short and often semicircular.
- May be attended by ants.

Aphidinae, Macrosiphoninae

**3.** Antennae usually longer than ¾ of body length, often longer than the body, on clearly visible antennal tubercles. If antennal tubercles are unclear, then reddish or green with black antennae. In dense colonies with ant visitation on *Tanacetum*…………………………………**(4)**

**3*.** Antennae ½ or 2/3 of body length, i.e., shorter than ¾ of body length. Mostly feeding along rims of leaves. Not attended by ants. Very small (adult 1.0-1.6 mm), yellowish or greenish globe-shaped. Legs, cornicles, and Cauda are almost colorless. The antennae are hard or, at most, slightly dark towards the tip only. Length of antennae. Cauda triangularly tongue-shaped, half as long as cornicles. Frontal sinus straight.…***Coloradoa tanacetina*** WALKER

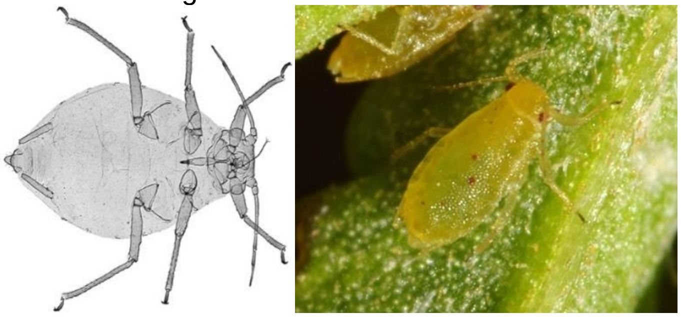

(InfluentialPoints, 2021)

*Coloradoa*

- Small green or reddish globe-shaped.
- Convex frons with no antennal tubercles.
- Antennae are always shorter than the body, and the terminal process is always longer than the base of the last antennal segment.
- Short dorsal body hairs and expanded at the tip.
- Apical rostral segment with concave sides.

**4.** Bright red to brownish red, feeding on the underside of bottom leaves, sometimes on shoots. Leaves often turn yellow on the upper side. Cornicles are long, thin, brown, and dark only at the tip and base. Antennae are a little shorter to (usually) longer than the body. Aptera body 2.5-3.4mm, pear-shaped. Legs yellowish, and black tibia's apices never with wax powder. Cauda is yellowish white (pale), relatively short, as long as the distance between bases of antennae. Antennal bases are divergent, not with frontal cusps directed inward or forward, i.e., inner sides not parallel to the longitudinal axis. Size: 2.5 to 3.4mm........................ ***Dactynotus tanaceti* (L.) (*Uroleucon tanaceti*)**

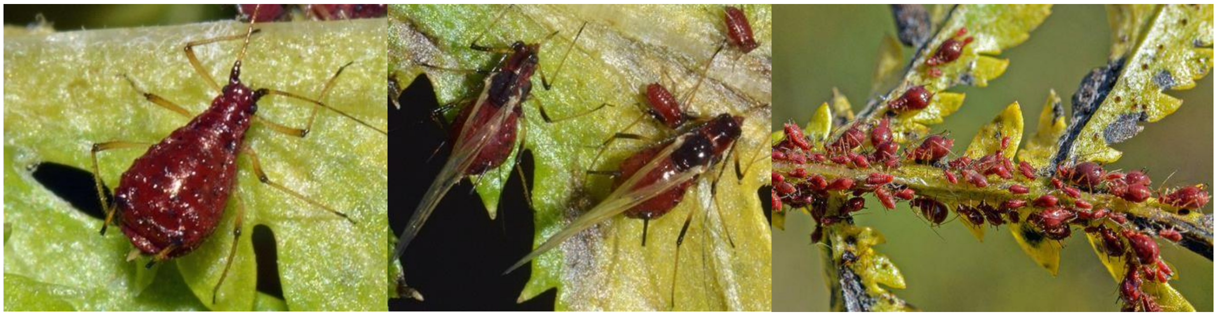

(InfluentialPoints, 2021)

*Uroleucon*

- Well-developed antennal tubercles with diverging sides.
- Antennae long as the body.
- Dorsal abdominal hairs are not capitated and are placed on sclerites, typically dark.

**4*.** Colored differently. If red or brown, then not very shiny or with wax powder…………… **(5)**

**5.** The body is completely covered with wax powder. Cornicles are primarily black, sometimes lighter, and then darker distally. Only at Asteraceae… ***Macrosiphoniella*** DEL GUERCIO **(6)**

**5*.** Roundish body, shiny or dull, at most the larvae weakly powdered with wax. Coloration brownish red. The abdomen is often green, with a large dark spot in the center. Mostly ant-tended. In inflorescences or upper stem parts with ant visitation Size: 2.2-2.8mm. Antennae as long as body length. Cornicles 1/5-1/6 of body length…….. ***Metopeurum fuscoviride* Stroyan**

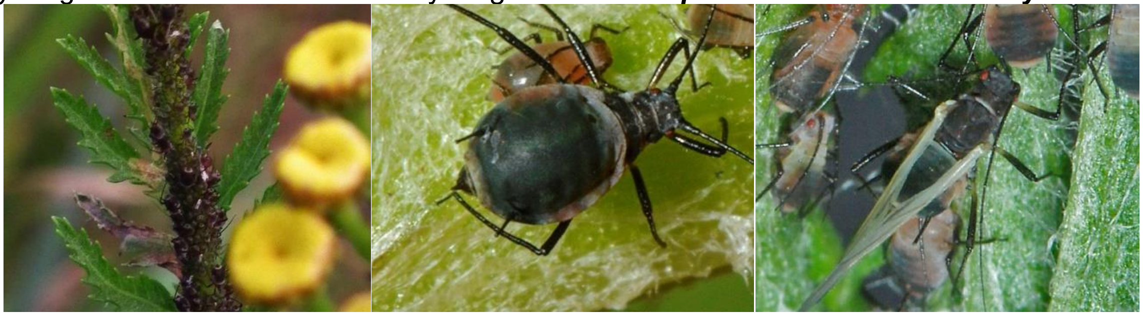

(InfluentialPoints, 2021)

*Metopeurum*

- Medium-sized aphids.
- Adult viviparous may be winged or wingless.
- Antennal tubercles are very weakly developed.
- Siphunculi dark over at least half of the length.
- Cauda tapering, triangular.

**6.** (rare on tansy) Tibiae and femora are black except for the basal part of the front femur, which is brown. Virgins green to yellowish green, sometimes reddish yellow in late summer (lighter green than *M. tanacetaria*). Dorsum has numerous hair-bearing black platelets (sclerites). Brown head. Medium to large (2.5-3.6mm), oval to elongate oval. White-grey waxy powder on the back, except for a spinal stripe on the abdomen and presiphuncular spots. Eyes mostly red. Antennae 1.0-1.2 times as long as the body with the terminal process is 3.7-4.3 times the length of the basal part. Si, cauda, F u B black except for the lighter thigh bases. Si 1/8-1/7 of body length, ¾-4/5 of caudal length. Mostly on stems, without ant visitation, some drop or play dead when disturbed. Pinkish-red male (middle picture)……… ***Macrosiphionella millefolii* (De Geer)**

Mostly on *Achillea millefolium*

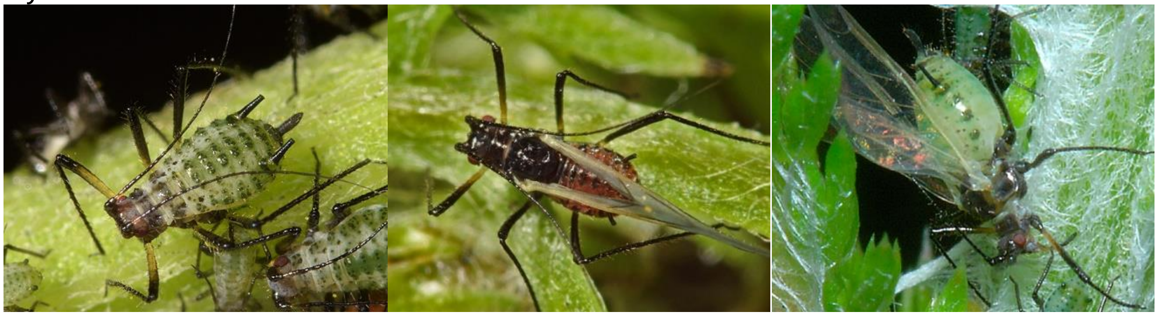

(InfluentialPoints, 2021)

**6*.** (common) Dorsum without black sclerites. Upper stem parts. Green or pinkish-brown, transversely powdered. The antennae, legs, siphunculi, and cauda are black. 3.2-3.9mm. Cornicles 1/7 of body length, 7/8 of caudal length. Antennae 1.0-1.3 times the body length, with the terminal process 2.9-3.5 times the length of the base. Siphunculi 0.1-0.2 times the body length and 0.6-0.9 times the length of the cauda. Colonies occur on the upper parts of the stem and between the flowers. Two color forms of *Macrosiphoniella tanacetaria* are green (more common) and pink. Not ant-tended (killed by ants)……………………***Macrosiphionella tanacetaria* (Kaltenbach)**

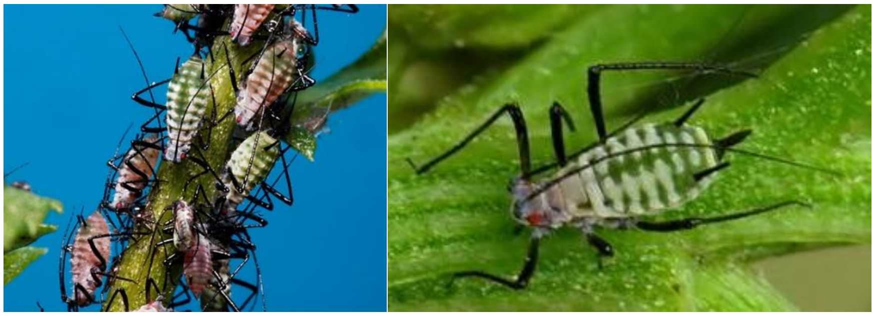

(InfluentialPoints, 2021)

*Macrosiphionella*

- Green to dark brown.
- Adult viviparous may be winged or wingless.
- Dorsum is not sclerotic; if pigmented, it is in small, localized hair-bearing sclerites.
- Siphunculi long, cylindrical, or with a slight taper from base to apex.
- No host alternation and no ant-attendance.

### GLOSSARY

**Antennal tubercles:** Base of antenna. Two lumps on the head, each of which bears one antenna. Sometimes large and protruding inwards (left picture).

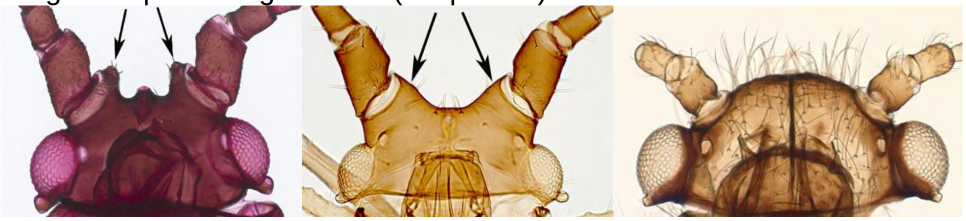

(parallel) (divergent) (absent)

(Favret & Miller, 2012)

**Antesiphuncular:** Immediately anterior to the base of each siphunculus.

**Cauda:** A taillike process or extension of the VIII abdominal segment.

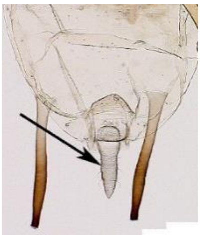

(Favret & Miller, 2012)

**Frons:** The middle part of the front of the head.

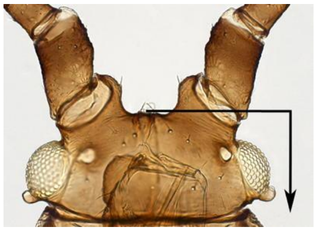

(Favret & Miller, 2012)

**Scleroites:** When the dorsal abdominal sclerotization consists of small sclerites at the base of the dorsal setae.

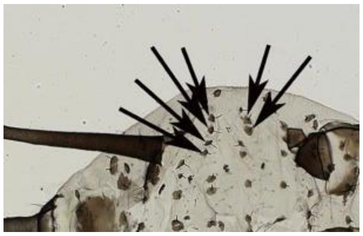

(Favret & Miller, 2012)

**Siphunculi/Cornicle:** Structures on the dorsum of the VI abdominal segment from which alarm pheromones are expelled.

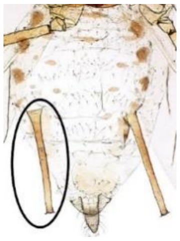

(Favret & Miller, 2012)

**Rostrum:** Beak-like labium usually of 4 segments, supporting the stylets.

**Terminal process:** The terminal antennal segment, usually the VI but sometimes the V segment, is divided into the base and the processus terminalis (PT).

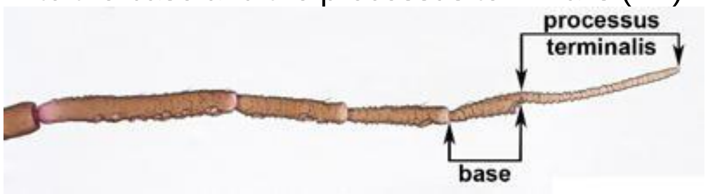

(Favret & Miller, 2012)

#### ***1.3 Statistical analysis***

*1.3.1 Effects of chemotype richness and chemotype presence on plot-level arthropod group occurrence*

Each arthropod group occurrence indicates whether an herbivore, flower visitor, predatory arthropod, or ant was found in a plot at the group level and lower taxonomic level when indicated. The occurrence was assessed at plot level for every year. Binomial GLMMs were fitted with CR as a fixed effect and block as a random effect to test the effect of CR. Separate analyses were conducted for each chemotype to investigate the effect of specific chemotypes on occurrence. In these models, chemotype presence (indicating whether a particular chemotype is present or absent in a plot) was used as a fixed effect, with block as a random effect in binomial GLMMs.

If chemotype presence had a significant effect on the response variable, we calculated the slope (e.g., correlation coefficient) of the linear relation between occurrence and CR.

*1.3.1.1 Statistical models*

*General models: Effect of chemotype richness on the occurrence*
GLMM: glmer(Occurrence of arthropod group or lower taxonomic level (when indicated) ~ CR + (1|Block), family = binomial)

*General models: Effect of chemotype presence on the occurrence*
GLMM: glmer(Occurrence of arthropod group or lower taxonomic level (when indicated) ~ CP + (1|Block), family = binomial)

*1.3.2 Effects of chemotype richness and chemotype presence on plot-level arthropod group abundance*

Herbivore abundance refers to the total number of aphids across all species accumulated in each plot over each year and to the cumulative plot-level number of individuals of all five aphid species separately across each year. Given the skewed nature of our data and the prevalence of zero counts in the herbivore abundance, we adopted Zero-Inflated Negative Binomial (ZINB) models when data transformations failed to meet LMM assumptions. We applied both models strategically based on the distribution of each subset of data. The goodness of fit was assessed through DHARMa plots alongside tests for overdispersion and zero inflation.

To assess the effects of CR on flower visitor abundance, we used the visitation rate (flower visitor abundance divided by the sum of the plot-level number of flower heads on the days of the respective surveys) as the response variable. This approach is supported as we found a significant influence of plant chemotype and CR on flower number and phenology (Ojeda-Prieto et al., 2024).

Predator refers to the total number of predators across all taxa/feeding guilds accumulated in each plot over the year and to the cumulative plot-level number of individuals of all predator categories separately across each year. Ant abundance refers to the total number of ants in each plot over the year.

To check the effect of CR on arthropod group abundance, LMMs were fitted with CR as a fixed effect and Block ID as a random effect on the total abundance of each arthropod group and lower taxonomic level.

To meet the model assumptions, we transformed the data. For herbivores, we squared-root the total abundance in 2021, 2022, and 2023, *A. fabae* abundance in 2021, *U. tanaceti* abundances in 2022 and 2023, and log-transformed *Me. fuscoviride* abundance in 2021. ZINB models were used for *A. fabae* abundances in 2022 and 2023, *B. cardui* abundances in the three years, *Ma. tanacetaria* abundances for 2021 and 2022, *Me. fuscoviride* abundance in 2022 and 2023, and *U. tanaceti* abundance in 2021. In the case of flower visitors, we used log-transformed flower visitor abundance across all orders and squared-transformed flower visitor abundance for each observed insect order without normalizing the number of flower heads. For predators, we squared-transformed the total number (plus one) of parasitized aphids in 2021, log-transformed the total number (plus one) of parasitized aphids in 2022, and used the untransformed predator abundances in 2023 (total predator count and by category).

We examined the effect of the presence/absence of a chemotype on each arthropod group abundance. In the case of herbivores, the same model structure, whether LMM or ZINB, was applied per chemotype, with chemotype presence (indicating whether a specific chemotype is present in the plot) and Block as fixed and random effects, respectively. We used LMMs for flower visitors, predators, and ants.

If chemotype presence had a significant effect on the abundance variable, we calculated the log-response ratio by taking the logarithm of the ratio of mean values for the “Presence” and “Absence” for each chemotype, allowing us to obtain the relative effect size for each chemotype, rather than absolute differences.

*1.3.2.1 Statistical models*

*General models:*

*Effect of chemotype richness on the abundance*

LMM: lmer(Abundance of arthropod group or lower taxonomic level (when indicated) ~ CR + (1|Block))

ZINB: glmmTMB(Abundance of arthropod group or lower taxonomic level (when indicated) ~ CR + (1|Block), zi = ~1, family ="nbinom2")

*Effect of chemotype presence on the abundance*

LMM: lmer(Abundance of arthropod group or lower taxonomic level (when indicated) ~ CP + (1|Block))

ZINB: glmmTMB(Abundance of arthropod group or lower taxonomic level (when indicated) ~ CP + (1|Block), zi = ~1, family ="nbinom2")

*1.3.3 Temporal effects*

Although a discussion of detailed temporal effects by sampling date is beyond the scope of this manuscript, we acknowledge that such temporal differences may occur. To visualize these temporal effects, we analyze CR's effect on the occurrence and abundance for each species, group, and order of herbivores, flower visitors, predators, and ants across the different sampling dates. Binomial GLMMs were applied to each sampling date occurrence with CR as the fixed factor and block as the random factor. Only when occurrence was higher than 60% (cut-off, otherwise indicated in grey in the table of Supplementary Information – S2) were abundance analyses made using LMMs. The choice of a 60% cut-off for occurrence when conducting abundance analyses is a strategic approach to ensure that LMM-based abundance analyses are conducted on a subset of data that aligns well with the model's assumptions. The cut-off is based on statistical considerations related to the distribution of data and the suitability of different modeling approaches.

We performed separate analyses for each sampling date, with abundance as the response variable, CR as the explanatory variable, and block as the random factor. Untransformed data were used for occurrence and abundance, except when indicated otherwise. For abundance, we squared root-transformed herbivore abundance for *total herbivore in 2021, A. fabae in 2021,* and *U. tanaceti in 2022,* and predator abundance in 2023 for *Araneae, Coleoptera, Diptera,* and *Parasitoid.* Log-transformation with an added constant of 1 was applied to the herbivore abundance for *total herbivores in 2022* and *2023, Ma. tanacetaria in 2022, U. tanaceti in 2023*, and flower visitor abundance in 2021 for the *visitation rate across all orders*. The slope corresponds to the correlation coefficient of the linear relationship between group occurrence or group abundance and CR.

### 2. Supplementary results

#### *2.1 Population patterns in the herbivore community*

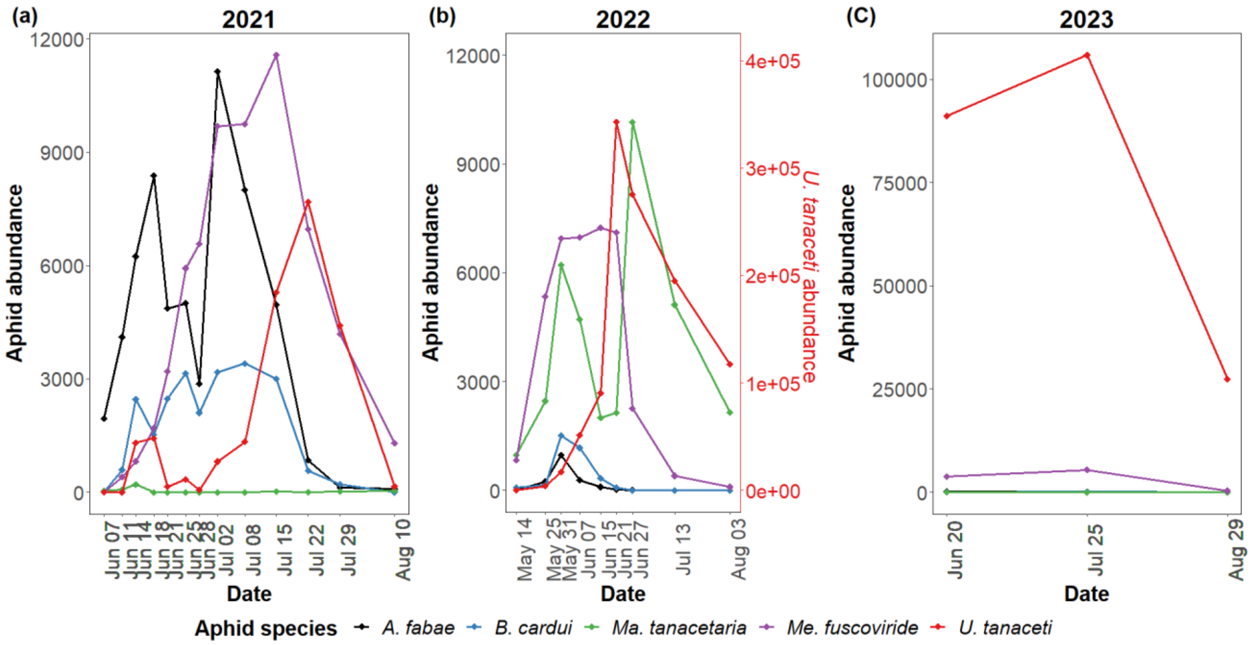

**Figure S1-3.** Total abundance of herbivores showing five different aphid species (*Aphis fabae, Brachycaudus cardui, Macrosiphoniella tanacetaria, Metopeurum fuscoviride,* and *Uroleucon tanaceti*) during the observation dates in 2021 **(a),** 2022 **(b)**, and 2023 **(c).** Each species is represented by a different color (black, blue, purple, green, and red). The points on each line represent the abundance of the corresponding aphid species on a specific date, while the lines illustrate the trend of aphid abundance over time. The dashed lines indicate the maximum abundance of each species throughout the year. The second y-axis in panel b is scaled differently for *U. tanaceti* species.

#### *2.2 Temporal effects*

Although an in-depth analysis by sampling dates will not be discussed here in detail, it is crucial to recognize temporal differences. Supplementary Information 2 (SI2) provides a detailed breakdown of the relationship between chemotype richness and the occurrence and abundance of various ecological groups across different sampling dates and years. For instance, most relationships with plot-level chemical richness in occurrence data were week, if significant. However, for abundance, these temporal patterns were sometimes very clear. Particularly for herbivore abundance, there was a positive relationship between plot-level chemodiversity in the

Weeks after establishing the field experiment in 2021, it changed towards a consistent negative correlation CR across all following sampling dates, particularly for specialist aphids. There was a positive correlation between flower visitor orders and ants. The effect on the abundance of different predator groups varied, but they tended to be positive when significant relationships were observed.

### *2.3 Supplementary tables*

**Table S1-2.** Summary of Binomial Linear Mixed-effect Models testing plot-level chemotype richness (CR) or chemotype identity (six different models, one for each chemotype: *Athu-Bthu, Bthu-high, Bthu-low, Chrys-acet, Mixed-high*, and *Mixed-low*) effects on herbivore occurrence, and Linear Mixed-effect Models (LMM) and Zero-Inflated Negative Binomial models (ZINB) testing herbivore abundance across all species (*All species*), as well as on each aphid species (*A. fabae, B. cardui, Ma. tanacetaria, Me. fuscoviride,* and *U. tanaceti*) in 2021. If all plots were occupied by at least one aphid in the year, we did not model occurrence (-), and this was the case for all species and *Me. fuscoviride*. Degrees of freedom, Wald's Chi-square statistics, and p-values are reported. Significant values (P <0.05) are reported in bold.

| | | | 2021 | | | | | |
|---|---|---|---|---|---|---|---|---|
| | | | **All species** | ***A. fabae*** | ***B. cardui*** | ***Ma. tanacetaria*** | ***Me. fuscoviride*** | ***U. tanaceti*** |
| **Response variable** | **Factor** | **d.f.** | **χ2 (p-value)** | **χ2 (p-value)** | **χ2 (p-value)** | **χ2 (p-value)** | **χ2 (p-value)** | **χ2 (p-value)** |
| Occurrence | CR | 1 | - | 1.04 (0.306) | 0.08 (0.784) | 0.62 (0.430) | - | 3.38 (0.066) |
| | Athu-Bthu | 1 | - | 0.55 (0.459) | 0.02 (0.890) | 0.15 (0.695) | - | 0.68 (0.409) |
| | Bthu-high | 1 | - | 0.31 (0.576) | 0.06 (0.800) | 1.34 (0.247) | - | 0.46 (0.495) |
| | Bthu-low | 1 | - | 0.57 (0.449) | 0.39 (0.531) | 0.35 (0.555) | - | 1.35 (0.245) |
| | Chrys-acet | 1 | - | 0.52 (0.470) | 2.48 (0.115) | 0.04 (0.845) | - | 0.51 (0.476) |
| | Mixed-high | 1 | - | 0.29 (0.587) | 0.25 (0.615) | 0.00 (1.000) | - | 7.47 **(0.006)** |
| | Mixed-low | 1 | - | 0.00 (0.998) | 0.01 (0.909) | 0.15 (0.695) | - | 0.42 (0.515) |
| Abundance | CR | 1 | 0.29 (0.588) | 0.03 (0.870) | 1.51 (0.219) | 1.13 (0.288) | 0.52 (0.471) | 0.38 (0.536) |
| | Athu-Bthu | 1 | 0.19 (0.665) | 0.05 (0.815) | 0.07 (0.793) | 1.21 (0.271) | 0.74 (0.390) | 3.60 (0.057) |
| | Bthu-high | 1 | 0.48 (0.487) | 0.07 (0.791) | 3.97 **(0.046)** | 0.00 (0.990) | 1.78 (0.182) | 5.98 **(0.014)** |
| | Bthu-low | 1 | 0.15 (0.699) | 0.09 (0.766) | 0.16 (0.688) | 0.05 (0.818) | 3.06 (0.080) | 0.14 (0.712) |
| | Chrys-acet | 1 | 0.15 (0.697) | 0.98 (0.322) | 6.16 **(0.013)** | 0.28 (0.595) | 0.27 (0.601) | 0.12 (0.725) |
| | Mixed-high | 1 | 2.24 (0.135) | 6.77 **(0.009)** | 0.01 (0.935) | 0.14 (0.705) | 0.03 (0.853) | 0.47 (0.491) |
| | Mixed-low | 1 | 0.82 (0.364) | 0.61 (0.434) | 0.00 (0.965) | 1.14 (0.286) | 0.35 (0.552) | 0.33 (0.565) |

**Table S1-3.** Summary of Binomial Linear Mixed-effect Models testing plot-level chemotype richness (CR) or chemotype identity (six different models, one for each chemotype: *Athu-Bthu, Bthu-high, Bthu-low, Chrys-acet, Mixed-high*, and *Mixed-low*) effects on herbivore occurrence, and Linear Mixed-effect Models (LMM) and Zero-Inflated Negative Binomial models (ZINB) testing herbivore abundance across all species (*All species*), as well as on each aphid species (*A. fabae, B. cardui, Ma. tanacetaria, Me. fuscoviride,* and *U. tanaceti*) in 2022. If all plots were occupied by at least one aphid in the year, we did not model occurrence (-); this was the case for all species and *U. tanaceti*. Degrees of freedom, Wald's Chi-square statistics, and p-values are reported. Significant values (P <0.05) are reported in bold.

| | | | **2022** | | | | | |
|---|---|---|---|---|---|---|---|---|
| | | | **All species** | ***A. fabae*** | ***B. cardui*** | ***Ma. tanacetaria*** | ***Me. fuscoviride*** | ***U. tanaceti*** |
| **Response variable** | **Factor** | **d.f.** | **χ2 (p-value)** | **χ2 (p-value)** | **χ2 (p-value)** | **χ2 (p-value)** | **χ2 (p-value)** | **χ2 (p-value)** |
| Occurrence | CR | 1 | - | 1.96 (0.162) | 1.43 (0.231) | 0.42 (0.515) | 0.97 (0.325) | - |
| | Athu-Bthu | 1 | - | 1.61 (0.204) | 0.26 (0.606) | 0.18 (0.669) | 0.06 (0.812) | - |
| | Bthu-high | 1 | - | 0.19 (0.662) | 4.16 **(0.041)** | 0.31 (0.576) | 1.66 (0.198) | - |
| | Bthu-low | 1 | - | 4.50 **(0.034)** | 6.42 **(0.011)** | 0.00 (0.953) | 1.26 (0.261) | - |
| | Chrys-acet | 1 | - | 0.00 (0.967) | 0.26 (0.606) | 0.52 (0.470) | 1.22 (0.270) | - |
| | Mixed-high | 1 | - | 6.68 **(0.010)** | 2.54 (0.111) | 0.40 (0.529) | 1.55 (0.213) | - |
| | Mixed-low | 1 | - | 4.89 **(0.027)** | 2.04 (0.153) | 0.14 (0.705) | 0.33 (0.568) | - |
| Abundance | CR | 1 | 7.63 **(0.006)** | 1.60 (0.206) | 2.57 (0.109) | 2.73 (0.098) | 1.10 (0.295) | 5.93 **(0.015)** |
| | Athu-Bthu | 1 | 3.46 (0.063) | 1.70 (0.192) | 0.88 (0.347) | 0.02 (0.882) | 0.08 (0.770) | 3.30 (0.069) |
| | Bthu-high | 1 | 0.01 (0.909) | 0.79 (0.374) | 0.74 (0.391) | 0.09 (0.760) | 0.73 (0.393) | 0.02 (0.898) |
| | Bthu-low | 1 | 0.26 (0.612) | 3.91 **(0.048)** | 3.39 (0.065) | 2.19 (0.139) | 5.85 **(0.016)** | 0.13 (0.717) |
| | Chrys-acet | 1 | 10.77 **(0.001)** | 0.48 (0.486) | 2.16 (0.142) | 0.29 (0.593) | 0.20 (0.654) | 12.77 **(<0.001)** |
| | Mixed-high | 1 | 4.81 **(0.028)** | 2.38 (0.122) | 1.01 (0.315) | 2.95 (0.086) | 0.32 (0.573) | 5.58 **(0.018)** |
| | Mixed-low | 1 | 0.18 (0.667) | 2.12 (0.146) | 0.00 (0.966) | 0.31 (0.575) | 0.53 (0.466) | 0.01 (0.938) |

**Table S1-4.** Summary of Binomial Linear Mixed-effect Models testing plot-level chemotype richness (CR) or chemotype identity (six different models, one for each chemotype: *Athu-Bthu, Bthu-high, Bthu-low, Chrys-acet, Mixed-high*, and *Mixed-low*) effects on herbivore occurrence, and Linear Mixed-effect Models (LMM) and Zero-Inflated Negative Binomial models (ZINB) testing herbivore abundance across all species (*All species*), as well as on each aphid species (*A. fabae, B. cardui, Ma. tanacetaria, Me. fuscoviride,* and *U. tanaceti*) in 2023. If all plots were occupied by at least one aphid in the year, we did not model occurrence (-); this was the case for all species and *U. tanaceti*. If no aphid was observed in the field, we did not model occurrence nor abundance (-); this was the case for *Ma. tanacetaria.* Degrees of freedom, Wald's Chi-square statistics, and p-values are reported. Significant values (P <0.05) are reported in bold.

| | | | **2023** | | | | | |
|---|---|---|---|---|---|---|---|---|
| **Response variable** | **Factor** | **d.f.** | **All species χ2 (p-value)** | ***A. fabae* χ2 (p-value)** | ***B. cardui* χ2 (p-value)** | ***Ma. tanacetaria* χ2 (p-value)** | ***Me. fuscoviride* χ2 (p-value)** | ***U. tanaceti* χ2 (p-value)** |
| Occurrence | CR | 1 | - | 0.16 (0.689) | 0.81 (0.369) | - | 9.24 **(0.002)** | - |
| | Athu-Bthu | 1 | - | 1.61 (0.204) | 1.22 (0.269) | - | 5.23 **(0.022)** | - |
| | Bthu-high | 1 | - | 0.19 (0.662) | 0.21 (0.648) | - | 1.58 (0.208) | - |
| | Bthu-low | 1 | - | 4.50 **(0.034)** | 1.10 (0.294) | - | 1.11 (0.740) | - |
| | Chrys-acet | 1 | - | 0.00 (0.967) | 1.23 (0.267) | - | 1.68 (0.194) | - |
| | Mixed-high | 1 | - | 6.68 **(0.010)** | 0.21 (0.649) | - | 9.42 **(0.002)** | - |
| | Mixed-low | 1 | - | 4.89 **(0.027)** | 0.46 (0.497) | - | 2.58 (0.108) | - |
| Abundance | CR | 1 | 4.39 **(0.036)** | 1.24 (0.265) | 5.56 **(0.018)** | - | 0.19 (0.663) | 4.44 **(0.035)** |
| | Athu-Bthu | 1 | 5.32 **(0.021)** | 7.39 **(0.006)** | 4.42 **(0.035)** | - | 0.50 (0.478) | 5.08 **(0.024)** |
| | Bthu-high | 1 | 0.09 (0.761) | 0.05 (0.818) | 40.71 **(<0.001)** | - | 0.21 (0.643) | 0.16 (0.688) |
| | Bthu-low | 1 | 0.06 (0.811) | 0.05 (0.818) | 0.778 (0.378) | - | 0.30 (0.586) | 0.13 (0.720) |
| | Chrys-acet | 1 | 2.77 (0.096) | 19.14 **(<0.001)** | 3.25 (0.071) | - | 1.48 (0.224) | 3.59 (0.058) |
| | Mixed-high | 1 | 0.65 (0.420) | 1.22 (0.270) | 79.26 **(<0.001)** | - | 0.56 (0.456) | 0.80 (0.372) |
| | Mixed-low | 1 | 3.96 **(0.046)** | 0.56 (0.453) | 0.10 (0.748) | - | 0.23 (0.631) | 3.92 **(0.048)** |

**Table S1-5.** Summary of Binomial Linear Mixed-effect Models testing plot-level chemotype richness (CR) or chemotype identity (six different models, one for each chemotype: *Athu-Bthu, Bthu-high, Bthu-low, Chrys-acet, Mixed-high*, and *Mixed-low*) effects on flower visitor occurrence, and Linear Mixed-effect Models (LMM) testing on visitation rate across all orders (*All orders*), as well as on each order (Coleoptera, Diptera, Hemiptera, Hymenoptera, and Lepidoptera) in 2021. If all plots were occupied by at least one pollinator, we did not model occurrence (-); this was the case for all *orders*. Visitation rate, calculated as the cumulative plot-level number of flower-visiting insects divided by the sum of the plot-level number of flower heads on the days of the flower-visitor surveys, is used as the response variable in the models to evaluate the effect of plot-level chemotype richness on the flower visitor abundances. Degrees of freedom, Wald's Chi-square statistics, and p-values are reported. Significant values ($P < 0.05$) are reported in bold.

| | | | 2021 | | | | | |
|---|---|---|---|---|---|---|---|---|
| | | | **All orders** | **Coleoptera** | **Diptera** | **Hemiptera** | **Hymenoptera** | **Lepidoptera** |
| **Response variable** | **Factor** | **d.f.** | **$\chi^2$ (p-value)** | **$\chi^2$ (p-value)** | **$\chi^2$ (p-value)** | **$\chi^2$ (p-value)** | **$\chi^2$ (p-value)** | **$\chi^2$ (p-value)** |
| Occurrence | CR | 1 | - | 0.46 (0.497) | 0.16 (0.688) | 0.79 (0.373) | 1.10 (0.294) | 3.87 **(0.049)** |
| | Athu-Bthu | 1 | - | 0.99 (0.319) | 0.43 (0.512) | 2.08 (0.149) | 0.28 (0.594) | 5.15 **(0.023)** |
| | Bthu-high | 1 | - | 0.26 (0.613) | 0.15 (0.634) | 0.24 (0.625) | 0.09 (0.763) | 0.44 (0.507) |
| | Bthu-low | 1 | - | 2.91 (0.088) | 0.62 (0.432) | 1.64 (0.201) | 5.30 **(0.021)** | 1.78 (0.152) |
| | Chrys-acet | 1 | - | 0.37 (0.544) | 3.73 (0.053) | 0.27 (0.602) | 0.46 (0.499) | 0.27 (0.604) |
| | Mixed-high | 1 | - | 1.63 (0.202) | 0.15 (0.694) | 2.15 (0.143) | 0.81 (0.369) | 0.05 (0.825) |
| | Mixed-low | 1 | - | 2.27 (0.132) | 0.018 (0.895) | 0.28 (0.594) | 0.00 (0.942) | 0.92 (0.336) |
| Abundance | CR | 1 | 10.71 **(0.001)** | 16.78 **(<0.001)** | 2.68 (0.102) | 0.54 (0.463) | 11.56 **(<0.001)** | 8.80 **(0.003)** |
| | Athu-Bthu | 1 | 1.11 (0.293) | 3.61 (0.057) | 1.10 (0.293) | 0.40 (0.526) | 1.78 (0.182) | 7.53 **(0.006)** |
| | Bthu-high | 1 | 3.18 (0.074) | 3.03 (0.082) | 0.71 (0.400) | 0.05 (0.826) | 1.99 (0.158) | 1.52 (0.217) |
| | Bthu-low | 1 | 0.51 (0.474) | 7.00 **(0.008)** | 0.02 (0.894) | 0.00 (0.993) | 7.58 **(0.006)** | 1.21 (0.272) |
| | Chrys-acet | 1 | 5.98 **(0.014)** | 6.38 **(0.012)** | 0.51 (0.475) | 0.61 (0.436) | 2.50 (0.113) | 1.65 (0.199) |
| | Mixed-high | 1 | 6.95 **(0.008)** | 0.98 (0.322) | 2.71 (0.100) | 1.58 (0.208) | 3.12 (0.077) | 0.36 (0.547) |
| | Mixed-low | 1 | 0.49 (0.484) | 2.95 (0.086) | 0.17 (0.679) | 0.06 (0.799) | 0.61 (0.434) | 2.39 (0.122) |

**Table S1-6.** Summary of Linear Mixed-effect Models (LMM) testing plot-level chemotype richness (CR) or chemotype identity (six different models, one for each chemotype: *Athu-Bthu, Bthu-high, Bthu-low, Chrys-acet, Mixed-high*, and *Mixed-low*) effects on flower visitor abundance across all orders (*All orders*), as well as on each order (Coleoptera, Diptera, Hemiptera, Hymenoptera, and Lepidoptera) in 2021. Degrees of freedom, Wald's Chi-square statistics, and p-values are reported. Significant values (P <0.05) are reported in bold.

| | | | **2021** | | | | | |
|---|---|---|---|---|---|---|---|---|
| **Response variable** | **Factor** | **d.f.** | **All orders**<br>**χ2 (p-value)** | **Coleoptera**<br>**χ2 (p-value)** | **Diptera**<br>**χ2 (p-value)** | **Hemiptera**<br>**χ2 (p-value)** | **Hymenoptera**<br>**χ2 (p-value)** | **Lepidoptera**<br>**χ2 (p-value)** |
| | CR | 1 | 43.66 **(<0.001)** | 21.90 **(<0.001)** | 3.83 (0.050) | 1.31 (0.252) | 15.21 **(<0.001)** | 7.40 **(0.006)** |
| | Athu-Bthu | 1 | 11.01 **(<0.001)** | 4.63 **(0.031)** | 1.99 (0.158) | 1.11 (0.293) | 2.39 (0.122) | 7.47 **(0.006)** |
| | Bthu-high | 1 | 8.26 **(0.004)** | 5.30 **(0.021)** | 0.85 (0.356) | 0.00 (0.939) | 2.47 (0.116) | 0.95 (0.330) |
| Abundance | Bthu-low | 1 | 31.04 **(<0.001)** | 20.44 **(<0.001)** | 3.26 (0.071) | 1.10 (0.293) | 18.88 **(<0.001)** | 2.29 (0.130) |
| | Chrys-acet | 1 | 0.16 (0.691) | 0.52 (0.473) | 0.98 (0.322) | 0.24 (0.628) | 0.38 (0.537) | 0.80 (0.372) |
| | Mixed-high | 1 | 7.19 **(0.007)** | 2.01 (0.156) | 2.68 (0.102) | 2.05 (0.153) | 3.78 (0.052) | 0.10 (0.751) |
| | Mixed-low | 1 | 5.66 **(0.017)** | 4.22 **(0.040)** | 0.95 (0.328) | 0.09 (0.761) | 1.22 (0.269) | 2.10 (0.148) |

**Table S1-7.** Summary of Binomial Linear Mixed-effect Models testing plot-level chemotype richness (CR) or chemotype identity (six different models, one for each chemotype: *Athu-Bthu, Bthu-high, Bthu-low, Chrys-acet, Mixed-high*, and *Mixed-low*) effects on predator occurrence, and Linear Mixed-effect Models (LMM) testing on predator abundance across all taxa/feeding guilds (*All*), as well as on each (Araneae, Coleoptera, Dermaptera, Diptera, and Parasitized aphid) in 2021. If all plots were occupied by at least one predator, we did not model occurrence (-); this was the case for all *groups*. We did not assess abundance except for parasitized aphids. Degrees of freedom, Wald's Chi-square statistics, and p-values are reported. Significant values (P <0.05) are reported in bold.

| | | | 2021 | | | | | |
|---|---|---|---|---|---|---|---|---|
| | | | **All** | **Araneae** | **Coleoptera** | **Dermaptera** | **Diptera** | **Parasitized aphid** |
| **Response variable** | **Factor** | **d.f.** | **$\chi^2$ (p-value)** | **$\chi^2$ (p-value)** | **$\chi^2$ (p-value)** | **$\chi^2$ (p-value)** | **$\chi^2$ (p-value)** | **$\chi^2$ (p-value)** |
| Occurrence | CR | 1 | - | 0.28 (0.597) | 3.22 (0.073) | 1.14 (0.285) | 1.05 (0.304) | 0.02 (0.894) |
| | Athu-Bthu | 1 | - | 0.00 (1.000) | 2.59 (0.107) | 0.04 (0.845) | 1.62 (0.203) | 0.30 (0.584) |
| | Bthu-high | 1 | - | 0.00 (0.999) | 0.00 (1.000) | 0.00 (1.000) | 0.40 (0.525) | 2.65 (0.103) |
| | Bthu-low | 1 | - | 0.00 (1.000) | 1.32 (0.252) | 0.09 (0.768) | 1.49 (0.223) | 0.17 (0.680) |
| | Chrys-acet | 1 | - | 0.00 (0.971) | 0.97 (0.324) | 0.00 (1.000) | 0.22 (0.636) | 2.82 (0.093) |
| | Mixed-high | 1 | - | 0.00 (1.000) | 0.00 (1.000) | 0.00 (1.000) | 1.22 (0.269) | 0.36 (0.551) |
| | Mixed-low | 1 | - | 0.00 (1.000) | 2.59 (0.107) | 0.04 (0.845) | 0.37 (0.544) | 4.71 **(0.030)** |
| Abundance | CR | 1 | - | - | - | - | - | 0.77 (0.381) |
| | Athu-Bthu | 1 | - | - | - | - | - | 0.62 (0.431) |
| | Bthu-high | 1 | - | - | - | - | - | 1.19 (0.275) |
| | Bthu-low | 1 | - | - | - | - | - | 0.10 (0.747) |
| | Chrys-acet | 1 | - | - | - | - | - | 6.30 **(0.012)** |
| | Mixed-high | 1 | - | - | - | - | - | 0.74 (0.391) |
| | Mixed-low | 1 | - | - | - | - | - | 1.75 (0.186) |

**Table S1-8.** Summary of Binomial Linear Mixed-effect Models testing plot-level chemotype richness (CR) or chemotype identity (six different models, one for each chemotype: *Athu-Bthu, Bthu-high, Bthu-low, Chrys-acet, Mixed-high*, and *Mixed-low*) effects on predator occurrence, and Linear Mixed-effect Models (LMM) testing on predator abundance across all taxa/feeding guilds (*All*), as well as on each (Araneae, Coleoptera, Diptera, Hemiptera, and Parasitized aphid) in 2022. If all plots were occupied by at least one predator, we did not model occurrence (-); this was the case for all taxa/feeding guilds. We did not assess abundance except for parasitized aphids. Degrees of freedom, Wald's Chi-square statistics, and p-values are reported. Significant values (P <0.05) are reported in bold.

| | | | 2022 | | | | | |
|---|---|---|---|---|---|---|---|---|
| **Response variable** | **Factor** | **d.f.** | **All**<br>**χ2 (p-value)** | **Araneae**<br>**χ2 (p-value)** | **Coleoptera**<br>**χ2 (p-value)** | **Diptera**<br>**χ2 (p-value)** | **Hemiptera**<br>**χ2 (p-value)** | **Parasitized aphid**<br>**χ2 (p-value)** |
| Occurrence | CR | 1 | - | 0.02 (0.884) | 0.00 (0.986) | 0.13 (0.716) | 6.54 **(0.010)** | 2.69 (0.101) |
| | Athu-Bthu | 1 | - | 0.00 (1.000) | 0.00 (1.000) | 0.03 (0.869) | 0.03 (0.866) | 0.12 (0.731) |
| | Bthu-high | 1 | - | 0.00 (0.999) | 0.00 (0.981) | 0.09 (0.767) | 0.66 (0.417) | 0.54 (0.464) |
| | Bthu-low | 1 | - | 0.00 (1.000) | 0.00 (1.000) | 0.02 (0.883) | 0.88 (0.350) | 2.47 (0.116) |
| | Chrys-acet | 1 | - | 0.00 (0.974) | 0.00 (1.000) | 1.02 (0.313) | 3.13 (0.077) | 5.55 **(0.018)** |
| | Mixed-high | 1 | - | 0.00 (0.999) | 0.00 (1.000) | 0.30 (0.582) | 3.13 (0.077) | 0.08 (0.784) |
| | Mixed-low | 1 | - | 0.00 (1.000) | 0.00 (1.000) | 0.09 (0.763) | 0.14 (0.707) | 0.03 (0.870) |
| Abundance | CR | 1 | - | - | - | - | - | 2.71 (0.100) |
| | Athu-Bthu | 1 | - | - | - | - | - | 1.01 (0.315) |
| | Bthu-high | 1 | - | - | - | - | - | 0.03 (0.859) |
| | Bthu-low | 1 | - | - | - | - | - | 2.87 (0.090) |
| | Chrys-acet | 1 | - | - | - | - | - | 0.87 (0.351) |
| | Mixed-high | 1 | - | - | - | - | - | 0.28 (0.598) |
| | Mixed-low | 1 | - | - | - | - | - | 0.32 (0.571) |

**Table S1-9.** Summary of Binomial Linear Mixed-effect Models testing plot-level chemotype richness (CR) or chemotype identity (six different models, one for each chemotype: *Athu-Bthu, Bthu-high, Bthu-low, Chrys-acet, Mixed-high*, and *Mixed-low*) effects on predator occurrence, and Linear Mixed-effect Models (LMM) testing on predator abundance across all taxa/feeding guild (*All*), as well as on each (Araneae, Coleoptera, Dermaptera, Diptera, Hemiptera, Parasitized aphid, Aphid parasitoid, and Other Hymenoptera) in 2023. If all plots were occupied by at least one predator, we did not model occurrence (-); this was the case for all taxa/feeding guilds and each of them. If we did not find a taxa/feeding guild in the field, we did not model occurrence nor abundance (-); this was the case for Parasitized aphids. Degrees of freedom, Wald's Chi-square statistics, and p-values are reported. Significant values (P <0.05) are reported in bold.

| | | | 2023 | | | | | | | | |
|---|---|---|---|---|---|---|---|---|---|---|---|
| | | | **All** | **Araneae** | **Coleoptera** | **Dermaptera** | **Diptera** | **Hemiptera** | **Parasitized aphid** | **Aphid parasitoid** | **Other Hymenoptera** |
| **Response variable** | **Factor** | **d.f.** | **χ2 (p-value)** | **χ2 (p-value)** | **χ2 (p-value)** | **χ2 (p-value)** | **χ2 (p-value)** | **χ2 (p-value)** | **χ2 (p-value)** | **χ2 (p-value)** | **χ2 (p-value)** |
| | CR | 1 | - | - | - | - | - | - | - | - | - |
| | Athu-Bthu | 1 | - | - | - | - | - | - | - | - | - |
| | Bthu-high | 1 | - | - | - | - | - | - | - | - | - |
| Occurrence | Bthu-low | 1 | - | - | - | - | - | - | - | - | - |
| | Chrys-acet | 1 | - | - | - | - | - | - | - | - | - |
| | Mixed-high | 1 | - | - | - | - | - | - | - | - | - |
| | Mixed-low | 1 | - | - | - | - | - | - | - | - | - |
| | CR | 1 | 0.06 (0.800) | 0.12 (0.725) | 0.38 (0.538) | 1.85 (0.174) | 1.91 (0.167) | 0.94 (0.332) | - | 2.38 (0.123) | 0.14 (0.706) |
| | Athu-Bthu | 1 | 0.87 (0.350) | 0.97 (0.324) | 0.03 (0.868) | 8.51 **(0.004)** | 0.22 (0.642) | 3.55 (0.060) | - | 0.38 (0.535) | 2.45 (0.117) |
| | Bthu-high | 1 | 0.02 (0.900) | 0.00 (0.951) | 0.21 (0.649) | 0.38 (0.539) | 0.05 (0.815) | 0.53 (0.465) | - | 0.92 (0.338) | 0.37 (0.540) |
| Abundance | Bthu-low | 1 | 2.20 (0.138) | 0.13 (0.720) | 2.50 (0.114) | 0.42 (0.517) | 1.75 (0.186) | 0.12 (0.730) | - | 0.19 (0.666) | 0.76 (0.383) |
| | Chrys-acet | 1 | 4.16 **(0.041)** | 0.47 (0.492) | 3.85 **(0.050)** | 2.95 (0.086) | 3.78 (0.052) | 0.52 (0.471) | - | 0.49 (0.484) | 0.05 (0.819) |
| | Mixed-high | 1 | 0.84 (0.361) | 1.62 (0.204) | 0.64 (0.425) | 0.20 (0.655) | 1.20 (0.273) | 0.02 (0.882) | - | 1.65 (0.198) | 0.41 (0.520) |
| | Mixed-low | 1 | 2.07 (0.150) | 0.12 (0.729) | 0.00 (0.989) | 1.43 (0.232) | 0.92 (0.336) | 2.39 (0.122) | - | 3.83 (0.050) | 0.07 (0.794) |

**Table S1-10.** Summary of Binomial Linear Mixed-effect Models testing plot-level chemotype richness (CR) or chemotype identity (six different models, one for each chemotype: *Athu-Bthu, Bthu-high, Bthu-low, Chrys-acet, Mixed-high*, and *Mixed-low*) effects on ant occurrence, and Linear Mixed-effect Models (LMM) testing on ant abundance in 2021, 2022, and 2023. If all plots were occupied at least by one ant in the year, we did not model occurrence (-); this was the case for 2023. We assessed abundance only in 2023. Degrees of freedom, Wald's Chi-square statistics, and p-values are reported. Significant values (P <0.05) are reported in bold.

| **Response variable: Ants** | | | **2021** | **2022** | **2023** |
|---|---|---|---|---|---|
| | **Factor** | **d.f.** | **χ2 (p-value)** | **χ2 (p-value)** | **χ2 (p-value)** |
| | CR | 1 | 0.00 (0.992) | 2.13 (0.144) | - |
| | Athu-Bthu | 1 | 0.00 (0.979) | 0.00 (0.948) | - |
| | Bthu-high | 1 | 0.00 (0.980) | 0.00 (1.000) | - |
| Occurrence | Bthu-low | 1 | 0.00 (0.978) | 0.04 (0.837) | - |
| | Chrys-acet | 1 | 0.00 (0.955) | 0.02 (0.891) | - |
| | Mixed-high | 1 | 0.00 (0.999) | 0.00 (0.957) | - |
| | Mixed-low | 1 | 0.00 (0.975) | 0.00 (0.946) | - |
| | CR | 1 | - | **-** | 3.15 (0.076) |
| | Athu-Bthu | 1 | - | **-** | 0.80 (0.370) |
| | Bthu-high | 1 | - | **-** | 1.77 (0.183) |
| Abundance | Bthu-low | 1 | - | **-** | 0.30 (0.584) |
| | Chrys-acet | 1 | - | **-** | 6.58 **(0.010)** |
| | Mixed-high | 1 | - | **-** | 0.33 (0.564) |
| | Mixed-low | 1 | - | **-** | 1.36 (0.244) |

**Table S1-11. Significant effects of plot-level chemotype presence on herbivore species.** Relative changes were calculated by the logarithm of the ratio of mean values for the "Presence" and "Absence" groups of each chemotype (columns) for each respective herbivore response variable (rows). The fill color of each tile represents the slope value for the corresponding group and date. The color scale ranges from blue (positive effect of chemotype presence) to white (no effect) to red (negative effect of chemotypes), with the midpoint set to 0; stronger colors represent stronger effects of the respective chemotype on plot-level response variables.

| | | | CHEMOTYPE | | | | | |
|---|---|---|---|---|---|---|---|---|
| | | | **Athu-Bthu** | **Bthu-high** | **Bthu-low** | **Chrys-acet** | **Mixed-high** | **Mixed-low** |
| **Occupancy** | 2021 | All species | | | | | | |
| | | *A. fabae* | | | | | | |
| | | *B. cardui* | | | | | | |
| | | *Ma. tanacetaria* | | | | | | |
| | | *Me. fuscoviride* | | | | | | |
| | | *U. tanaceti* | | | | | **-0.288** | |
| | 2022 | All aphid species | | | | | | |
| | | *A. fabae* | | | **0.488** | | **0.619** | **-0.540** |
| | | *B. cardui* | | **0.330** | **-0.433** | | | |
| | | *Ma. tanacetaria* | | | | | | |
| | | *Me. fuscoviride* | | | | | | |
| | | *U. tanaceti* | | | | | | |
| | 2023 | All aphid species | | | | | | |
| | | *A. fabae* | | | **-0.629** | | **0.916** | **-0.097** |
| | | *B. cardui* | | | | | | |
| | | *Me. fuscoviride* | **-0.410** | | | | **-0.601** | |
| | | *U. tanaceti* | | | | | | |
| **Abundance** | 2021 | All species | | | | | | |
| | | *A. fabae* | | | | | **0.297** | |
| | | *B. cardui* | | **0.449** | | **0.663** | | |
| | | *Ma. tanacetaria* | | | | | | |
| | | *Me. fuscoviride* | | | | | | |
| | | *U. tanaceti* | | **0.815** | | | | |
| | 2022 | All aphid species | | | | **-0.046** | **-0.034** | |
| | | *A. fabae* | | | **-0.481** | | | |
| | | *B. cardui* | | | | | | |
| | | *Ma. tanacetaria* | | | | | | |
| | | *Me. fuscoviride* | | | **-0.341** | | | |
| | | *U. tanaceti* | | | | **-0.050** | **-0.037** | |
| | 2023 | All aphid species | **-0.134** | | | | | **-0.134** |
| | | *A. fabae* | **-1.916** | | | **-0.689** | | |
| | | *B. cardui* | **-2.096** | **-0.498** | | | **-0.269** | |
| | | *Me. fuscoviride* | | | | | | |
| | | *U. tanaceti* | **-0.133** | | | | | **-0.131** |

**Table S1-12. Significant effects of plot-level chemotype presence on flower visitor orders.** Relative changes were calculated by the logarithm of the ratio of mean values for the "Presence" and "Absence" groups of each chemotype (columns) for each respective flower visitor response variable (rows). The fill color of each tile represents the slope value for the corresponding group and date. The color scale ranges from blue (positive effect of chemotype presence) to white (no effect) to red (negative effect of chemotypes), with the midpoint set to 0; stronger colors represent stronger effects of the respective chemotype on plot-level response variables.

| | | | CHEMOTYPE | | | | | |
|---|---|---|---|---|---|---|---|---|
| | | | **Athu-Bthu** | **Bthu-high** | **Bthu-low** | **Chrys-acet** | **Mixed-high** | **Mixed-low** |
| **Occupancy** | 2021 | All orders | | | | | | |
| | | Coleoptera | | | | | | |
| | | Diptera | | | | | | |
| | | Hemiptera | | | | | | |
| | | Hymenoptera | | | **0.487** | | | |
| | | Lepidoptera | **0.730** | | | | | |
| **Abundance** | 2021 | All orders | | | | **0.440** | **0.315** | |
| | | Coleoptera | | | **0.251** | **0.256** | | |
| | | Diptera | | | | | | |
| | | Hemiptera | | | | | | |
| | | Hymenoptera | | | **0.332** | | | |
| | | Lepidoptera | **0.748** | | | | | |

**Table S1-13. Significant effects of plot-level chemotype presence on predatory groups.** Relative changes were calculated by the logarithm of the ratio of mean values for the “Presence” and “Absence” groups of each chemotype (columns) for each respective predatory response variable (rows). The fill color of each tile represents the slope value for the corresponding group and date. The color scale ranges from blue (positive effect of chemotype presence) to white (no effect) to red (negative effect of chemotypes), with the midpoint set to 0; stronger colors represent stronger effects of the respective chemotype on plot-level response variables.

| | | | **CHEMOTYPE** | | | | | |
|---|---|---|---|---|---|---|---|---|
| | | | **Athu-Bthu** | **Bthu-high** | **Bthu-low** | **Chrys-acet** | **Mixed-high** | **Mixed-low** |
| **Occurrence** | 2021 | All | | | | | | |
| | | Araneae | | | | | | |
| | | Coleoptera | | | | | | |
| | | Dermaptera | | | | | | |
| | | Diptera | | | | | | |
| | | Parasitized aphid | | | | | | **-0.254** |
| | 2022 | All | | | | | | |
| | | Araneae | | | | | | |
| | | Coleoptera | | | | | | |
| | | Diptera | | | | | | |
| | | Hemiptera | | | | | | |
| | | Parasitized aphid | | | | **-0.279** | | |
| | 2023 | All | | | | | | |
| | | Araneae | | | | | | |
| | | Coleoptera | | | | | | |
| | | Dermaptera | | | | | | |
| | | Diptera | | | | | | |
| | | Hemiptera | | | | | | |
| | | Aphid parasitoid | | | | | | |
| | | Other Hymenoptera | | | | | | |
| **Abundance** | 2021 | Parasitized aphid | | | | **0.432** | | |
| | 2022 | Parasitized aphid | | | | | | |
| | 2023 | All orders | | | | **-0.039** | | |
| | | Araneae | | | | | | |
| | | Coleoptera | | | | **-0.034** | | |
| | | Dermaptera | **0.093** | | | | | |
| | | Diptera | | | | | | |
| | | Hemiptera | | | | | | |
| | | Parasitized aphid | | | | | | |
| | | Aphid parasitoid | | | | | | |
| | | Other Hymenoptera | | | | | | |

**Table S1-14. Significant effects of plot-level chemotype presence on ants.** Relative changes were calculated by the logarithm of the ratio of mean values for the "Presence" and "Absence" groups of each chemotype (columns) for each respective ant response variable (rows). The fill color of each tile represents the slope value for the corresponding group and date. The color scale ranges from blue (positive effect of chemotype presence) to white (no effect) to red (negative effect of chemotypes), with the midpoint set to 0; stronger colors represent stronger effects of the respective chemotype on plot-level response variables.

| | | CHEMOTYPE | | | | | |
|---|---|---|---|---|---|---|---|
| | **Response variable: Ants** | **Athu-Bthu** | **Bthu-high** | **Bthu-low** | **Chrys-acet** | **Mixed-high** | **Mixed-low** |
| **Occurrence** | 2021 | | | | | | |
| | 2022 | | | | | | |
| | 2023 | | | | | | |
| **Abundance** | 2023 | | | | **0.056** | | |
